# Molecular Basis for Maternal Inheritance of Human Mitochondrial DNA

**DOI:** 10.1101/2023.05.12.540615

**Authors:** William Lee, Angelica Zamudio-Ochoa, Gina Buchel, Petar Podlesniy, Nuria Marti Gutierrez, Margalida Puigros, Anna Calderon, Hsin-Yao Tang, Li Li, Amy Koski, Ramon Trullas, Shoukhrat Mitalipov, Dmitry Temiakov

## Abstract

Uniparental inheritance of mitochondrial DNA (mtDNA) is an evolutionary trait found in nearly all eukaryotes. In many species, including humans, the sperm mitochondria are introduced to the oocyte during fertilization^1, 2^. The mechanisms hypothesized to prevent paternal mtDNA transmission include ubiquitination of the sperm mitochondria and mitophagy^3, 4^. However, whether these mechanisms play a decisive role in paternal mtDNA elimination has been disputed^5, 6^. We found that mitochondria in human spermatozoa are devoid of mtDNA and lack mitochondrial transcription factor A (TFAM), the major nucleoid protein required to protect, maintain, and transcribe mtDNA. During spermatogenesis, sperm cells express an isoform of TFAM, which retains the mitochondrial pre-sequence, ordinarily removed upon mitochondrial import. Phosphorylation of this pre-sequence prevents mitochondrial import and directs TFAM to the spermatozoon nucleus. TFAM re-localization from the mitochondria of spermatogonia to the spermatozoa nucleus directly correlates with the elimination of mitochondrial DNA, thereby explaining maternal inheritance in this species.

---

Uniparental inheritance of mtDNA is one of evolution’s few enduring pillars. Except for some fungi and mussels, most eukaryotes inherit their mtDNA from a single parent, most commonly a mother. The uniparental mtDNA inheritance pattern evolved repeatedly, and a multitude of mechanisms developed to exclude the paternal (sperm) mitochondria from a zygote^7^. This suggests the existence of selection pressure, as losing the uniparental inheritance was weakening and, perhaps, even detrimental for some species^8^. However, the benefits of uniparental inheritance for fitness are not immediately clear. Intuitively, having genetically divergent copies of mtDNA may complicate their cooperation with the nuclear genome and generate “selfish” mtDNA molecules, which replicate efficiently but cannot support adequate energy production^9^. Indeed, heteroplasmy, or the presence of different haplotypes of mtDNA, results in genetic instability and adverse physiological effect in mice^10^.

Despite the overwhelming dominance of uniparental inheritance, the underlying molecular mechanisms that prevent biparental inheritance are not understood. Downregulation of mtDNA during spermatogenesis occurs in fruit flies and mice^11, 12^. It has also been reported that ubiquitination of the sperm mitochondria leads to their targeted degradation by the proteasome-dependent proteolytic machinery after fertilization in mammals^3^. Selective degradation of the paternal mitochondria by autophagy in the embryonic cytoplasm has been documented in nematodes^4^. However, experiments show a lack of mitophagy in mammalian oocytes^5, 6^, and thus the mechanisms of maternal inheritance of human mtDNA remain obscure.

Previous studies reported a large variation of mtDNA copy number in human spermatozoa - from 1 to 1000 genomes per cell^13–15^. To accurately measure the mtDNA copy number in human sperm cells, we used Droplet Digital PCR (ddPCR), which allows a determination of the absolute number of DNA molecules in a cell (**Fig. 1A**). Because the method does not require isolation of the genomic material, and allows quantification of nuclear and mtDNA genomes in the same sample with high analytical sensitivity and a limit of detection below a single genome copy per microliter of sample, ddPCR is ideally suited for measuring low copy number nucleic acids^16^. Amplification targeted two single-copy genes in the nuclear genome (TBP and TEFM) and two mitochondrial genes – CYTB and ND1 (**Fig. 1B,C**). We found that the sperm cells contain, on average, 0.58 copies of mtDNA (**Fig. 1D**). Each spermatozoon contains 50-70 mitochondria, corresponding to less than 0.01 mtDNA molecules per mitochondrion. While extremely low, this number likely accounts for the background mtDNA detected in a few contaminating cells (namely leucocytes), which can harbor up to 100 mtDNA per cell in their mitochondria. Using an orthogonal strategy, we performed *in situ* hybridization of mtDNA in human testicular tissue using RNAScope (**Fig. 1E**). This method allows the detection of single DNA molecules per cell. An intense signal corresponding to mtDNA was detected in the mitochondria of spermatogonia, undifferentiated germ cells at the periphery of the seminiferous tubules (**Fig. 1E**). The primary spermatocytes, the cells developed from spermatogonium during the second stage of spermatocytogenesis, showed coarse staining of mtDNA, indicating its reduction during sperm maturation. Finally, mature spermatozoa had no detectible mtDNA signal in the midpiece region, in agreement with our ddPCR analysis (**Fig. 1E**). We conclude that the mature *human spermatozoa are essentially devoid of mtDNA*, consistent with maternal inheritance of the mitochondrial genome in mammals.

**Figure 1.**
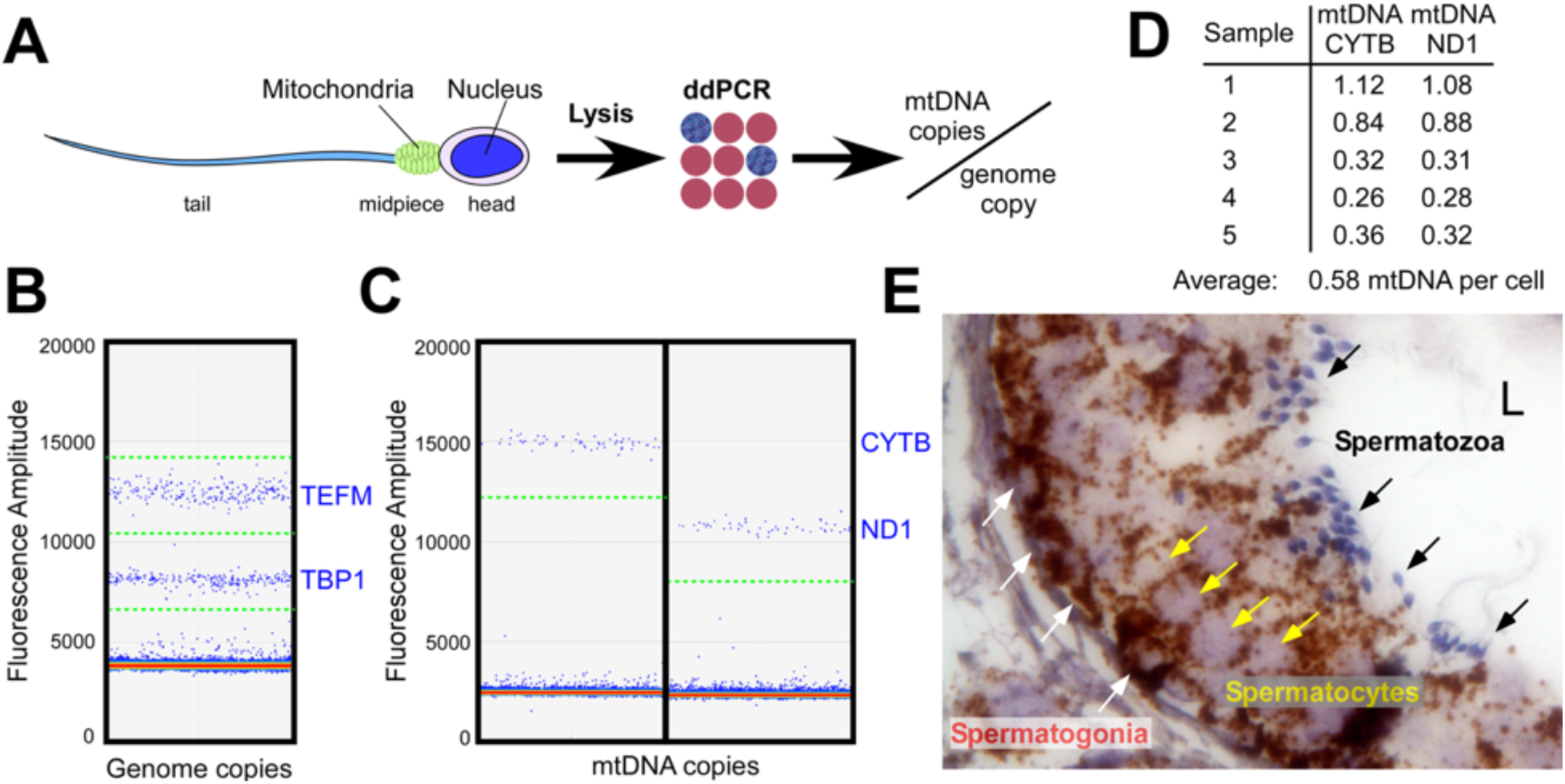
Mitochondria of human spermatozoa contain no mtDNA. **A.** Schematics of the mtDNA copy number measurements in sperm cells by ddPCR. **B, C.** Representative 1D droplet scatter plots of ddPCR analysis of spermatozoa samples. The multiplex analysis of amplification of two single-copy nuclear genes, TATA-box binding protein 1 and TEFM (B), and of two mitochondrial genes, cytochrome b, and ND1 (C) in the same spermatozoa sample is presented. Blue dots above the thresholds (dotted green lines) indicate droplets positive for the target amplicons. **D.** Summary data indicating mtDNA copy number per genome. **E**. *In situ* RNAScope labeling of mtDNA in developing sperm cells in seminiferous tubules of testicular tissue. MtDNA- brown, white arrows - spermatogonia, yellow – spermatocytes, black – spermatozoa. L – seminiferous tubule lumen.

We next looked at the presence of key proteins involved in transcription and replication of mtDNA in the mitochondria of human spermatozoa. We found that amounts of human mitochondrial RNA polymerase (POLRMT), the catalytic subunit of DNA polymerase PolG, and transcription elongation factor, TEFM, were below the detection limit of Western blotting (**Extended Data Fig. 1A**). More importantly, these proteins were absent from all published complete spermatozoa proteomes^17–19^. These findings demonstrate that the spermatozoa mitochondria cannot maintain, replicate or transcribe mtDNA. Unexpectedly, we found that human and monkey spermatozoa contain large amounts of the major mitochondrial nucleoid protein, transcription factor TFAM (**Fig. 2**).

**Figure 2.**
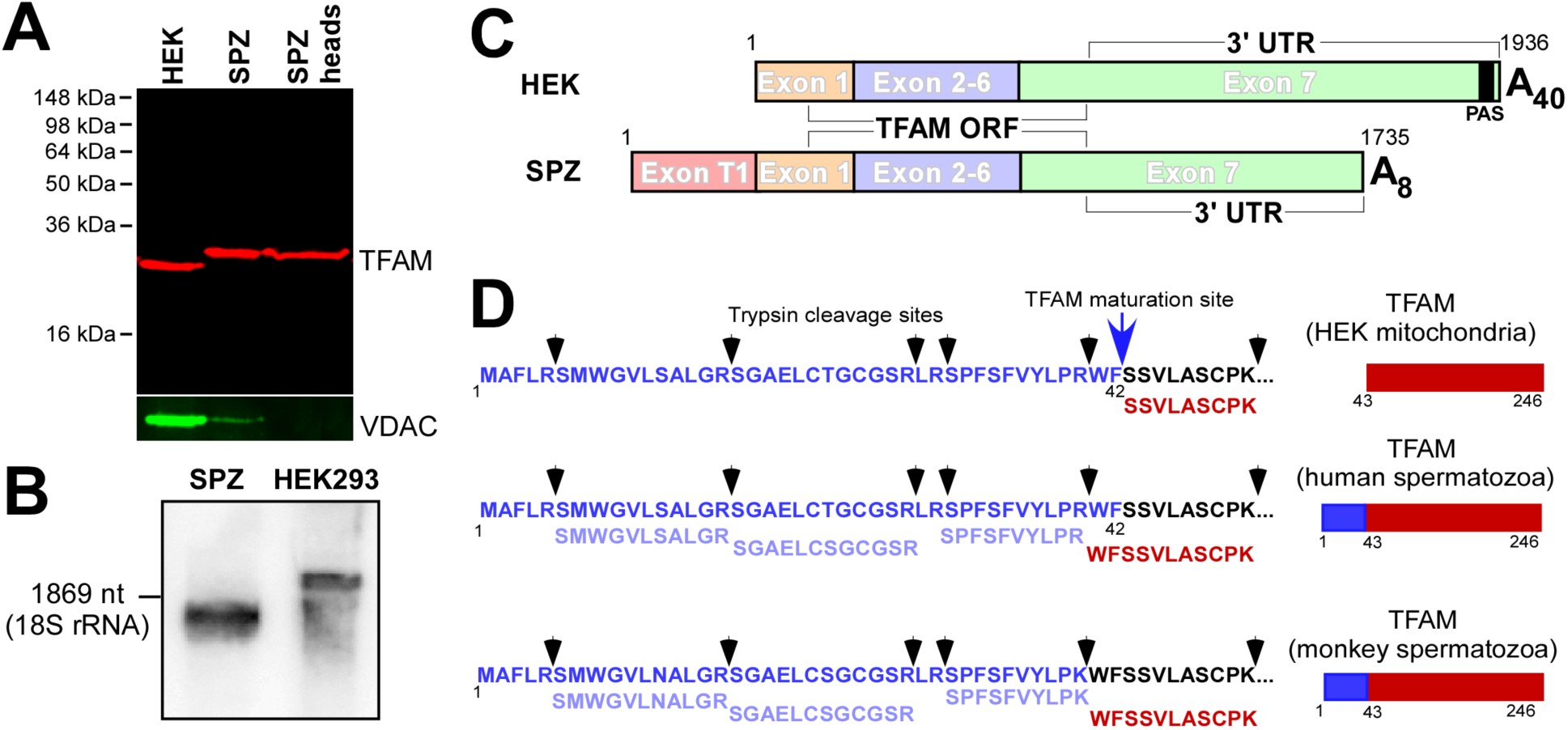
Mature sperm cells contain a specific isoform of TFAM. **A.** Western blot of mitochondrial lysate of HEK cells (lane 1), spermatozoa (lane 2), and spermatozoa in which the tails/midpieces have been removed (lane 3) are shown. Bottom - Western blot of the same gel stained with the anti-VDAC antibody. **B.** Sperm isoform of TFAM is a product of alternative splicing of the TFAM pre-mRNA. Northern blot of mRNA extracted from sperm or HEK cells. **C**. Schematic illustration of TFAM mRNA structure in somatic and sperm cells. PAS – polyadenylation signal. **D.** Sperm TFAM isoform contains a mitochondrial pre-sequence as revealed by LC-MS/MS data.

Western blot assays revealed a peculiar form of TFAM, which was ∼ 5 kDa larger than the mature (mitochondrial) TFAM from HEK cells (**Fig. 2A**). A previous study detected alternative splicing of TFAM mRNA during the spermatogonia maturation^20^. Indeed, we found that the mature sperm cells contain an alternatively spliced TFAM mRNA, not found in somatic cells (**Fig. 2B**). We mapped the 3’ UTR of this transcript and found that it is shorter than the one in somatic cells and contains an abbreviated, ∼ 8 nt long poly A-tail (**Fig. 2C, Extended Data Fig. 1**). In agreement with the previous data obtained for testis tissue, as the result of an alternative splicing event, the sperm TFAM mRNA features an additional exon T1 (**Fig. 2B**). However, the T1 exon does not alter the protein open reading frame, and thus the translation of the sperm mRNA isoform results in the synthesis of a full-size TFAM precursor (residues 1-246), as confirmed by LC-MS/MS analysis (**Fig. 2D, Extended Data Fig. 2**). Both human and Rhesus monkey sperm TFAM contain the peptides found in the mitochondrial targeting sequence (residues 1-42) but lack a partial tryptic fragment (evidence of protein maturation), suggesting that unlike the mitochondrial TFAM in somatic cells, this protein isoform does not undergo mitochondrial processing, in which this signal would be removed (**Fig. 2D**).

TFAM localizes exclusively to the mitochondrial reticulum in somatic cells (**Fig. 3A**). In contrast, confocal microscopy reveals TFAM in the head of spermatozoa but not in the mitochondria at the midpiece region (**Fig. 3B**). This is consistent with the presence of an unprocessed mitochondrial targeting sequence in the sperm TFAM isoform, as established by LC-MS/MS analysis. Since TFAM is absolutely required for mtDNA maintenance, replication, and transcription, and the reduction of its expression is linked to mtDNA elimination in all animal models tested^21–23^, the lack of TFAM in the spermatozoa mitochondria explains why these cells do not possess mtDNA.

**Figure 3.**
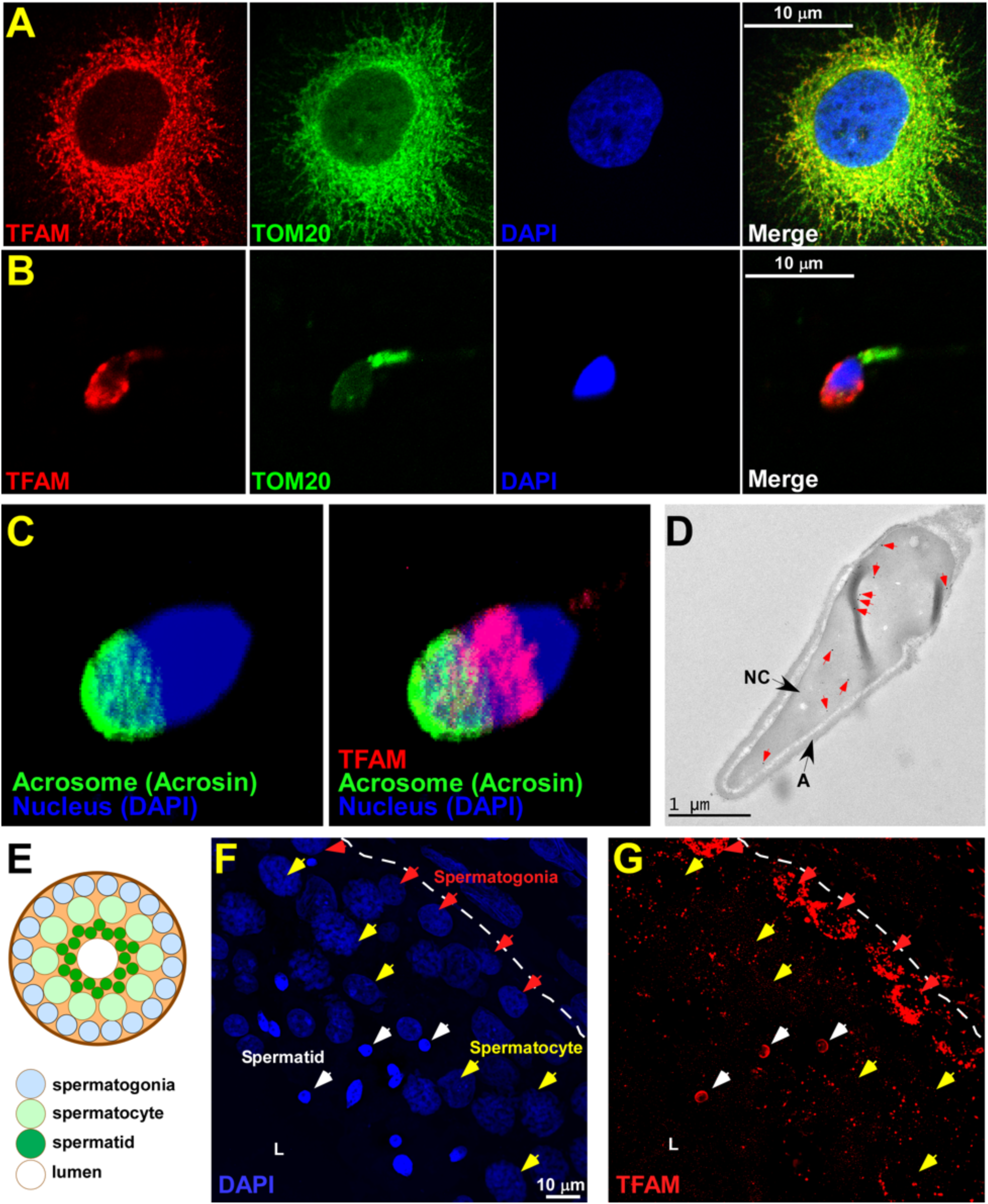
Mature sperm cells contain TFAM in the nucleus but not in the mitochondria. **A, B.** Confocal microscopy shows the localization of the endogenous TFAM (red) in mitochondria of HeLa cells (**A**) and the head of spermatozoa (**B**). Mitochondria staining with an anti-TOM20 antibody (green), and nucleus - DAPI (blue). **C.** Deconvolved 3D reconstructed confocal Z-stack image of spermatozoa. Staining with the anti-acrosin antibody (left) or anti-acrosin and anti-TFAM antibody (right). **D**. Cryo immunogold electron microscopy of spermatozoa cells reveals nuclear localization of TFAM. The head region is shown. Red arrows indicate gold particles. A - acrosome, NC- nucleus. **E.** Schematics of a transversal cut of a seminiferous tubule. **F, G** Human testicular tissue staining using DAPI (blue, G) and an anti-TFAM antibody (red, H). Red arrows - spermatogonia, yellow – spermatocytes, white – spermatids. The basement membrane is indicated by a dashed line. L – seminiferous tubule lumen.

Confocal microscopy revealed a nuclear localization of TFAM in the anterior half of the spermatozoa head, in proximity to a sperm-specific organelle, acrosome (**Fig. 3B)**. However, a detailed analysis of TFAM localization using 3D reconstruction of the deconvolved images suggests a clear co-localization of TFAM with the spermatozoon nucleus and not with the acrosome (**Fig. 3C**). Further, immunogold labeling of TFAM revealed the presence of gold-stained TFAM in the nucleus of the sperm cells (**Fig. 3D**), but not in the sperm mitochondria, as evident from the analysis of the images taken (n=25) (**Extended Data Fig. 3A,B**).

Our data suggest that during sperm cell development, TFAM should switch its localization from mitochondria to the nucleus. We stained human testicular tissue with an anti-TFAM antibody (**Fig. 3E-G**). An intense TFAM signal was detected in the mitochondria of spermatogonia (**Fig. 3G**). In contrast, primary spermatocytes showed only traces of TFAM, which partially overlaps with TOM20, indicative of mitochondrial localization (**Fig. 3F,G and Extended Data Fig. 3C**). Closer to the lumen, the immature sperm cells -round spermatids – reveal intense DAPI staining of their compacted nuclei, which co-localized with the TFAM signal (**Fig. 3F,G**). We speculate that sperm TFAM may already be present in the nuclei of the primary spermatocytes but becomes visible only when “concentrated” by the nuclear condensation in spermatids. Remarkably, the re-location of TFAM from the mitochondria of immature sperm cells to the spermatozoa nucleus coincides with the disappearance of mtDNA observed during spermatogenesis (**Fig. 1E**).

Cytosolic translation of some yeast mitochondrial proteins is mediated by PUF proteins that recognize the 3’ UTRs sequences and direct the mRNAs to the ribosomes associated with the mitochondrial outer membrane to enable a co-translational import ^24^. Assuming that a similar targeting system might function in other eukaryotes and considering that an alternatively spliced variant of TFAM mRNA has been found in sperm cells, we set out to investigate how the presence of the sperm-specific UTRs affects the trafficking of TFAM in spermatozoa. Transduction of HeLa cells with lentiviral constructs containing TFAM mRNA with either the sperm-specific (nuclear) or somatic (mitochondrial) 5’- and 3’- UTRs, or the mRNA lacking the UTRs altogether resulted in TFAM localization exclusively into the mitochondria, as evident by confocal microscopy (**Fig. 4A, Extended Data Fig. 3D,E**). In contrast, when the mitochondrial pre-sequence has been deleted, was TFAM localized to the cell nucleus (**Fig. 4B**). We speculate that the nuclear localization of TFAM lacking mitochondrial pre-sequence results from a predicted nearly-consensus bipartite nuclear localization signal in TFAM^25^.

**Figure 4.**
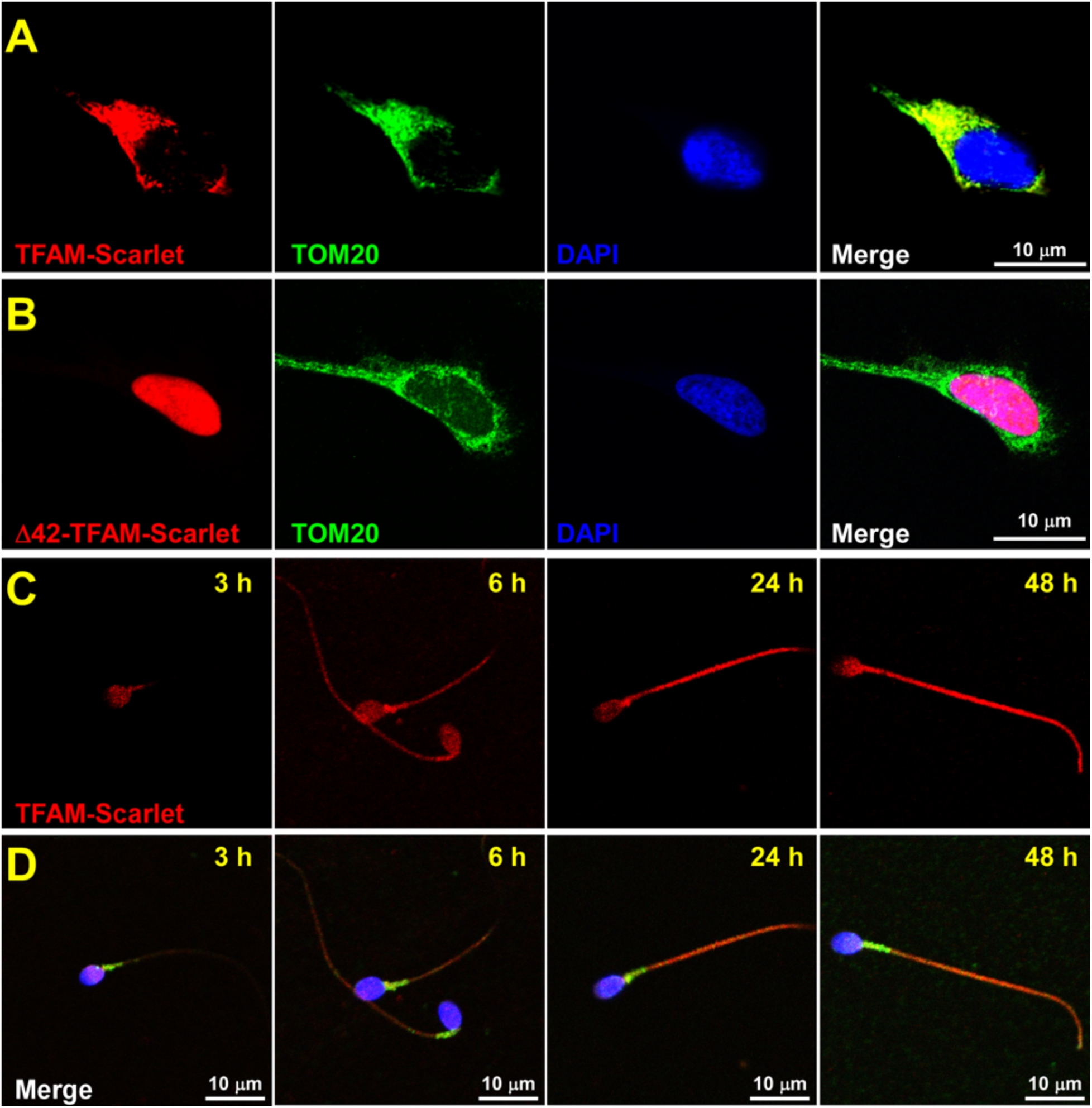
MTS, but not UTR of TFAM mRNA plays a role in its localization. **A.** UTR regions are not essential for TFAM localization in somatic cells. HeLa cells were transduced with the TFAM-mScarlet fusion having the nuclear 3’ and 5’ UTRs. **B.** Only a pre-sequence region is required for mitochondrial localization of TFAM in somatic cells. HeLa cells were transduced with TFAM-mScarlet fusion lacking the first 42 amino acids. **C, D.** Over-expressed TFAM-mScarlet fusion protein is found in the cytoplasm of sperm cells.

When investigating TFAM trafficking in sperm cells, we observed an efficient accumulation of TFAM-mScarlet in the cytoplasm, visible by a progression of the red fluorescent signal from the sperm head to its tail (**Fig. 4C)**. Interestingly, we did not detect TFAM accumulation in the nucleus or mitochondria (**Fig. 4D**). TFAM was localized in the cytoplasm of the sperm cells irrespective of the presence or absence of UTRs (**Extended Data Fig. 4A**). The extremely high packing density of the genomic material, about 10-fold higher than in somatic cells^26^, could explain the absence of exogenous TFAM in the sperm nucleus. Indeed, overexpression of the sperm nuclear protein, histone H2B, shows cytoplasmic localization of this protein in sperm cells but nuclear localization in somatic cells (**Extended Data Fig. 4B,C**). Notwithstanding, the import of proteins into sperm mitochondria appears active, as we detected mitochondrial localization of the overexpressed TOM20 in the transduced spermatozoa (**Extended Data Fig. 4D,E**). Most importantly, the lack of TFAM-mScarlet fluorescence signal in mitochondria of the transduced sperm cells suggests that the mitochondrial import of the over-expressed protein is being actively prevented.

To understand the reason behind the lack of TFAM import into the spermatozoa mitochondria, we transduced these cells with lentivirus carrying red fluorescent protein mScarlet fused to the mitochondrial pre-sequence of TFAM or of mtRNAP (**Fig. 5A,B and Extended Data Fig. 5**). While we observed mScarlet localization in sperm mitochondria when mtRNAP pre-sequence was used (**Extended Data Fig. 5A**), we detected no fluorescence signal inside mitochondria in the case of the TFAM pre-sequence (**Fig. 5A**), hinting at the key role of this region in sperm TFAM localization. Indeed, when ι142-TFAM-mScarlet fusion with the mtRNAP pre-sequence was used, this protein was localized to spermatozoa mitochondria (**Fig. 5B**).

**Figure 5.**
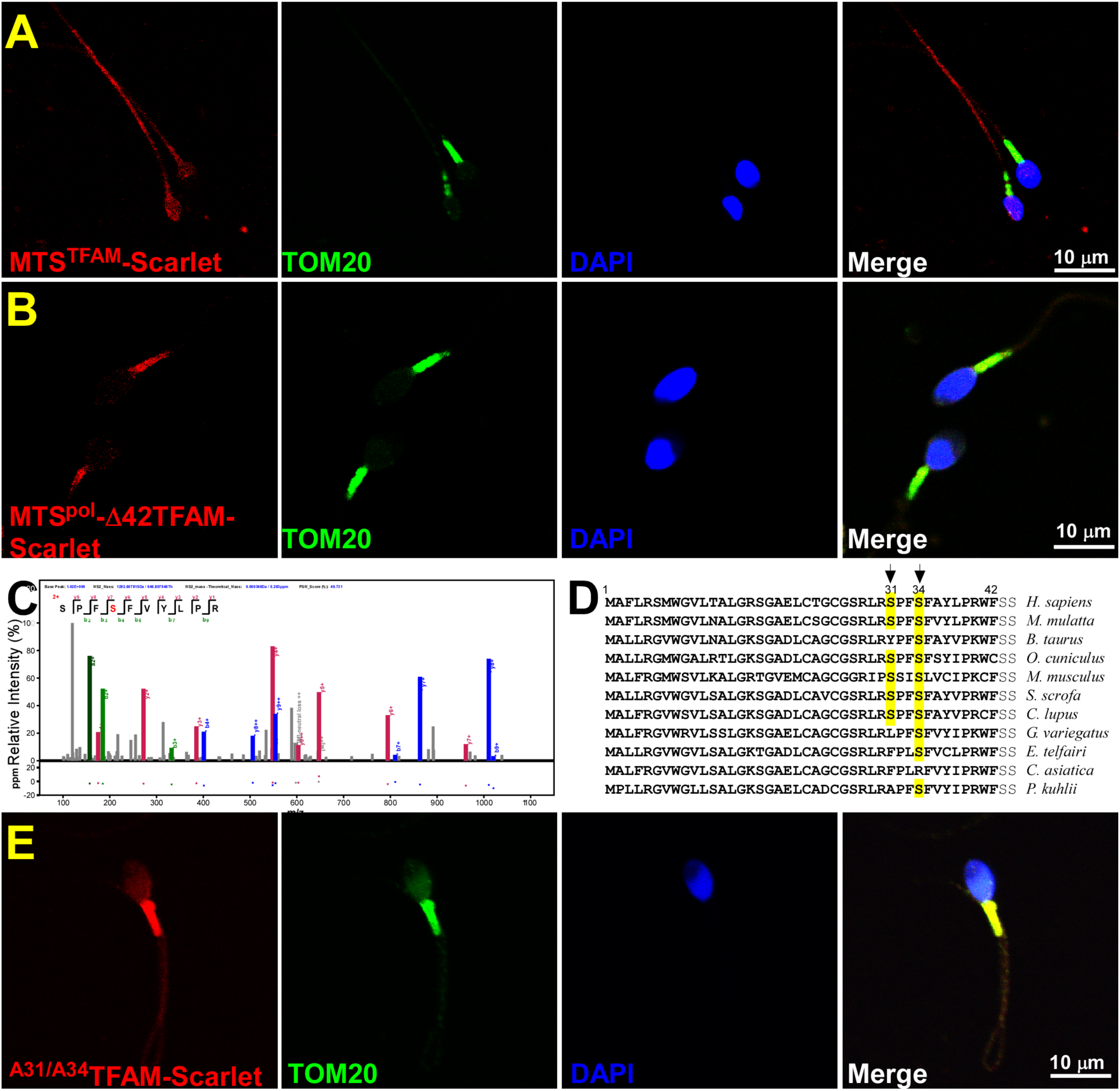
TFAM import to mitochondria is prevented in spermatozoa. **A.** MTS^TFAM^-mScarlet protein is localized to the sperm cytoplasm. **B.** MTS^pol^-1′42TFAM-mScarlet is localized to the sperm mitochondria. **C.** S34 residue in the sperm TFAM pre-sequence is phosphorylated. The MS/MS spectrum confirming the presence of Ser^34^ in TFAM is shown. m/z, mass-to-charge ratio. **D.** Conservation of the serine residues implicated in TFAM phosphorylation in mammalian species. **E.** TFAM^S31A/S34A^-mScarlet protein is localized to the sperm mitochondria.

Discovering the crucial role of the pre-sequence in preventing the mitochondrial import of TFAM prompted us to search for posttranslational modifications in this region. LC-MS/MS analysis of human sperm TFAM identified phosphorylation of the residue S34 in the pre-sequence (**Fig. 5C**). Analysis of the human sperm phosphoproteome confirmed the presence of phosphorylation at S34 and, in addition, at S31 in the sperm TFAM pre-sequence^27^. Mitochondrial pre-sequence recognition is sequence-independent and based on its secondary structure (commonly an α helix) and the positive charge of this region^28^. Phosphorylation of S31 and S34, conserved in mammalian TFAM (**Fig. 5D**), is expected to bring a large negative net charge to the pre-sequence region and prevent mitochondrial import.

While the function of MTS phosphorylation is largely unknown, several reports indicate inhibition of protein translocation into mitochondria^29–31^. To confirm that the phosphorylation of the TFAM pre-sequence region plays a role in preventing mitochondrial import of TFAM in spermatozoa, we substituted S31 and S34 residues with alanines and transduced sperm cells with lentivirus carrying this variant. Using confocal microscopy, we found that TFAM^S31A/S34A^ was localized exclusively to the sperm mitochondria (**Fig. 5E**).

## Concluding remarks

Maternal inheritance of mtDNA is a major paradigm that guides the existence and evolution of the vast majority of species; however, the molecular basis of this phenomenon and its benefits has remained unclear. Previous studies of human mitochondrial inheritance focused on post-fertilization mechanisms of elimination of paternal organelles and mtDNA from oocytes^3, 32^, while pre-fertilization mechanisms have never been reported. Here we demonstrate that phosphorylation-dependent re-localization of TFAM from the mitochondria to the nucleus of sperm cells during spermatogenesis results in mtDNA elimination and explains its maternal inheritance (**Fig. 6**).

**Figure 6.**
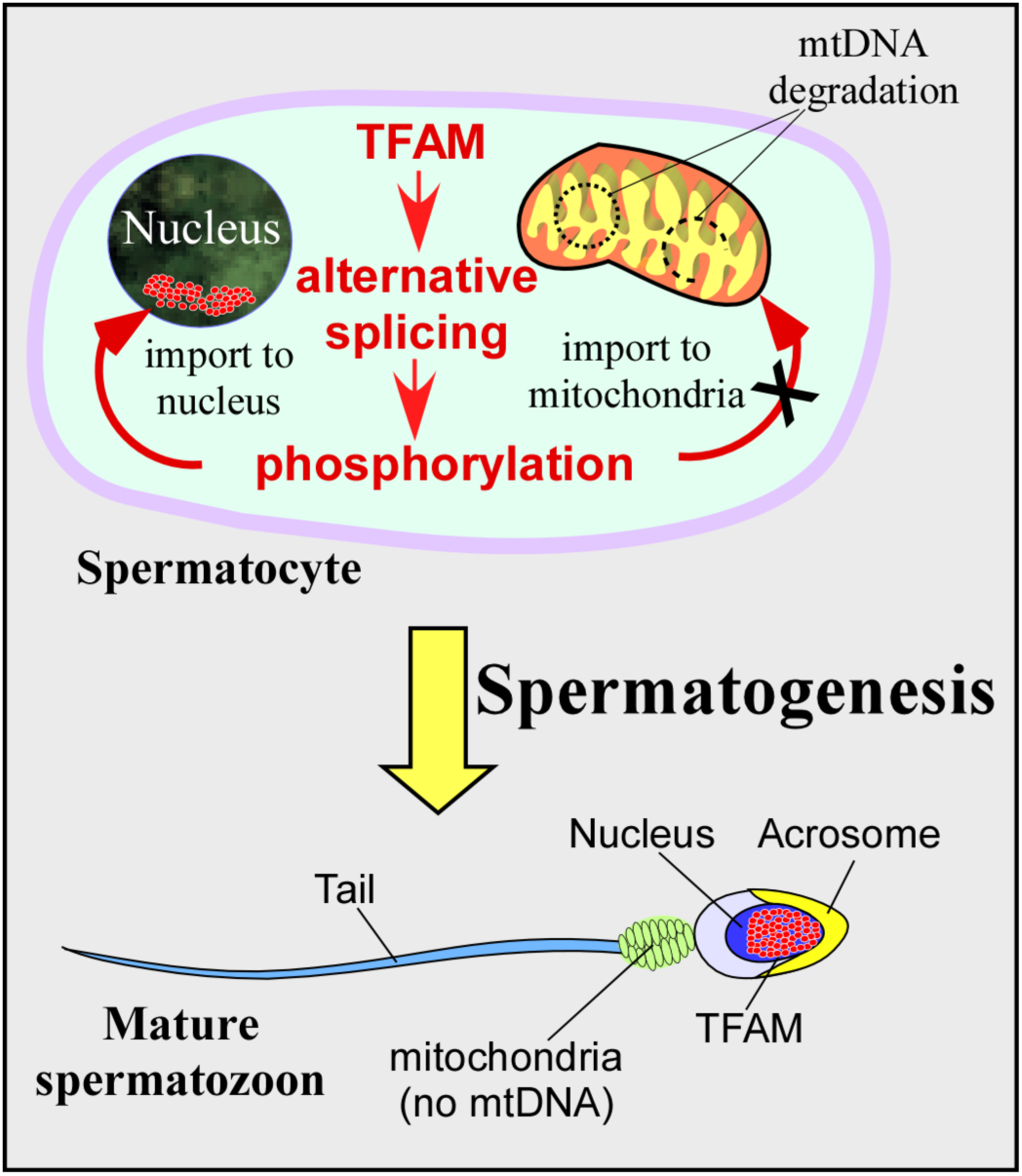
Sperm TFAM re-localization during spermatogenesis. TFAM redistribution during sperm maturation. During spermatogenesis, TFAM is phosphorylated and prevented from being imported to mitochondria, resulting in mtDNA degradation. Instead, TFAM is accumulated in the sperm nucleus.

The phenomenon of TFAM re-localization has important implications for the fields of human fertility and germ cell therapy^33^. Spermatozoa that are deficient in nuclear TFAM could account for unexplained male infertility. Indeed, elevated mtDNA levels were found in the sperm of men with severe oligoasthenospermia^15^. Our findings suggest that the sperm isoform of TFAM can serve as a novel biomarker of male infertility and may have clinical implications.

## Methods

### Cell culture

HEK293, HEK293T, and HeLa cells were cultured in DMEM containing 10% (v/v) FBS and 1% (v/v) Penicillin/Streptomycin (all from Gibco) at 37 °C and 5% CO_2_ in a humidified atmosphere. Adherent cells were harvested by incubation with 0.25% trypsin-EDTA. Cells were passaged and maintained at a 2.5 × 10^5^ cells/ml density.

### Gamete collection and analysis

Human subjects donations and study protocols were approved by Oregon Health & Science University (OHSU) IRB. Prior to gamete donations, informed consent was obtained from all human subjects on study-specific consent forms. Oversight defining research on human gametes and preimplantation embryos at OHSU undergoes strict regulatory review. All sample collections and sample preparations for analysis were performed at OHSU.

Semen collection was obtained via at-home semen collection kits or at OHSU Fertility Clinic. Participants were instructed to abstain from ejaculation for 24 h prior to collection. The ejaculate was maintained at body temperature until processing. Sperm samples were obtained from healthy adult donors and liquefied for 30-60 min at room temperature (RT). After liquefaction, the semen was loaded on AllGrad 90% (Cooper Surgical, #AG90-100) gradient solution and centrifuged at 1500 rpm for 15 min. The pellet containing spermatozoa was then rinsed in Quinn’s sperm washing medium (Cooper Surgical, #ART-1006) and centrifuged again at 1100 rpm for 6 min. The cells were counted using a hemocytometer and observed for the presence of the contaminating cells.

All animal experiments were performed following NIH Guidelines for Animal Research and approved by OHSU IACUC. All mice were maintained under specific pathogen-free and controlled lighting conditions (12 hours light/dark cycles) at OHSU. Six to eight weeks old BDF1 females were superovulated with 5 international units of pregnant mare’s serum gonadotropin (PMSG) and human chorionic gonadotropin (hCG). MII oocytes were collected from excised oviducts 14 h post hCG injection. Sperm from the adult Rhesus macaque (*Macaca mulatta*) was collected as described previously ^34^.

### Measurement of the absolute copy number of mtDNA

Purified human sperm cells (4,000,000 cells in 2 μl) were mixed with 8 μl of the pre-warmed lysis solution containing 10 M Urea and 30 mM DTT and incubated for 5 min at 95°C. Upon incubation, the lysed cells (5 μl) were transferred into a 1.5 ml tube containing 1 ml of DNA/RNA/protein solubilization reagent 100ST (DireCtQuant), mixed by vortexing, and incubated for 10 min at 95°C to complete the lysis and to dilute the sample to the optimal concentration of Urea (less than 50 mM). The resulting sample containing 2,000 lysed spermatozoa cells per μl of solution was used to measure the absolute copy number of mtDNA by Droplet Digital PCR (ddPCR). MtDNA was amplified using two primer pairs, mt64-ND1, and mt92-CYTB, which target two opposite regions of the mtDNA genome, as described^35^.

The droplet reaction consisted of 1x QX200™ ddPCR™ EvaGreen Supermix (1864033, Bio-Rad), a 0.1 μl aliquot of the target sample, and TBP73 (95 nM) and TEFM88 (140 nM) primers for genome analysis, or mt64-ND1(100 nM) and mt92-CYTB (135 nM) primers for mtDNA analyses. A restriction enzyme digestion was performed prior to partitioning into droplets by adding the Fast Digest enzymes HaeIII and MseI (0.5 U each) into the ddPCR reaction for nuclear genome, and AluI (1 U) for mtDNA amplification. The reactions were incubated for 15 min at 37° C. Non-template controls were included in each analysis plate to monitor possible reaction contamination. The PCR amplification was performed using the C1000 Touch Thermal Cycler (Bio-Rad) and the following thermal profile: 95° C 5 min; (95° C 30 sec; 60° C 1 min) 40 repeats; 4° C 5 min; 90° C 10 min.

The data analysis was performed with QuantaSoft Analysis Pro v1.0 using thresholds to distinguish single and double-positive droplet populations. The number of genomes was calculated by averaging the DNA copy number obtained using TBP73 and TEFM88. To determine the mtDNA amount per haploid genome in a sperm cell, the number of mtDNA copies obtained with the mt64-ND1 or the mt92-CYTB amplicon was divided by the number of genomes measured in the same sperm sample. Simple or multiplex analyses were performed using the QX200 Digital Droplet PCR platform (Bio-Rad Laboratories).

### Chromogenic In Situ Hybridization Assay (CISH) using human testicular tissue

The study involving testicular tissues was approved by the Thomas Jefferson University Research IRB protocol. Human testicular tissues from patients with testicular cancer were provided by the Department of Pathology at Thomas Jefferson University Hospital. Non-neoplastic human testicular parenchyma from the orchiectomy specimens was obtained. The tissues were sectioned at 4 μm in thickness, away from the tumor, and mounted onto slides. The tissues were fixed with 10% formalin for 24 h, immersed in a series of various concentrations of ethanol (70, 80, 95, 100%) and xylene followed by embedding into paraffin blocks with paraffin wax. Hematoxylin and eosin (H&E) staining has been performed. The H&E slides of testicular tissue were reviewed by a pathologist. The sections with normal spermatogenesis were marked, and adequate unstained sections were obtained from the formalin-fixed paraffin-embedded (FFPE) blocks for CISH and IHC.

CISH with mtDNA in testicular tissues was performed exactly as described in the manufacturer’s protocol (Advanced Cell Diagnostic), using RNAscope 2.5 HD Assay BROWN Detection Kit and Hs-MT-COX1-sense probe (#478051; ACD). The signal was developed by incubation with DAB (3,3′-diaminobenzidine) for 10 minutes at RT and counterstained with hematoxylin (Sigma), followed by a wash in 0.02% ammonia water before mounting. Digital Images were obtained using Olympus BX60 microscope at 20X magnification and the cellSens imaging software (Olympus).

### Mitochondria isolation

HeLa cells were gently homogenized with 25 strokes using a Teflon homogenizer (Thomas Scientifics) in 7.5 ml of HES buffer (20 mM HEPES, pH7.2, 0.25 M sucrose, and 0.1 mM EDTA), supplemented with 1 mM DTT, 0.1 mg/ml BSA, and 0.1 mM PMSF. Homogenates were centrifuged at 2,000 g for 5 minutes at 4 °C. Supernatants were collected and centrifuged again at 10,000 g for 12 minutes. The crude mitochondria were resuspended in HES buffer and further purified by a sucrose step gradient separation (15/23/34/60% sucrose in 20 mM HEPES, pH 7.2, 0.1 mM EDTA) at 100,000 g at 4 °C for 1 h. The purified mitochondria were resuspended in 100 μl of HES buffer, aliquoted, and stored at −20 °C.

### RNA purification

To extract the RNA from human spermatozoa (∼80 million cells in 100 μl), they were homogenized by vortexing with 100 mg of 0.2 mm stainless steel beads (Next Advance) in Lysis Buffer (Thermo Scientific), supplemented with guanidine thiocyanate and 2% 2-mercaptoethanol, for 5 minutes at RT. The RNA was isolated from the lysate using the GeneJET RNA Purification Kit (Thermo Scientific), aliquoted, and stored at −80^0^ C. To extract the RNA from HEK293 cells, they were lysed in 1 ml of TRI Reagent (Invitrogen) for 5 minutes at RT. 1-Bromo-3-chloropropane (100 μl) was added, vortexed for 15 seconds, and incubated for 10 minutes at RT. The lysate was then centrifuged at 12,000 g for 15 minutes at 4° C to separate the RNA-containing aqueous phase. The RNA was precipitated by mixing the aqueous phase with isopropanol and centrifuged at 12,000 g for 15 minutes at 4° C. The RNA pellet was washed with 75% ethanol and air-dried before resuspending in DEPC-treated water. The purified RNA was aliquoted and stored at −80^0^ C.

### Mapping of the 3’ UTR region of the mitochondrial and sperm isoform of TFAM

RNA samples were first treated with dsDNAse (Thermo Fisher Scientific) to remove any DNA contamination. The 3’ ends of the sperm and HEK293 TFAM mRNA species were mapped by 3’ RLM-RACE using the FirstChoice RLM-RACE Kit (Ambion) with modifications. The RNA adaptor provided by the kit was phosphorylated on its 5’ end using T4 Polynucleotide Kinase (NEB). RNA (1 μg) was ligated to the 5’-phosphorylated RNA adaptor using T4 RNA Ligase (Ambion). Following ligation, the TFAM mRNA was amplified by RT-PCR using the SuperScript IV One-Step RT-PCR System (Thermo Fisher Scientific). The RT-PCR reaction was further amplified by PCR using inner primers (**Extended Table 1**).

### LC-MS/MS analysis

To reduce the complexity of the sample for LC-MS/MS analysis, the band representing sperm TFAM was excised from 10% PAGE. Spermatozoa cells (∼ 5 million) were lysed in SDS sample buffer by sonication. Recombinant human TFAM was used as a protein marker to excise the band representing the sperm TFAM isoform. The excised protein band was washed and subjected to trypsin digestion. The gel bands were treated with tris(2-carboxyethyl)phosphine, alkylated with iodoacetamide, and the protein digested with trypsin. Tryptic digests were analyzed by LC-MS/MS using a Q Exactive Plus or Q Exactive HF mass spectrometer (ThermoFisher Scientific) coupled with a Nano-ACQUITY UPLC system (Waters) as previously described ^36, 37^. Peptides and proteins were identified using MaxQuant ^38^. MS/MS spectra were searched against the UniProt human or rhesus macaque protein database and a common contaminant database using full tryptic specificity with up to two missed cleavages and static carbamidomethylation of Cys. Variable modifications included in the search were oxidation of Met, deamidation of Asn, and protein N-terminal acetylation. To identify the TFAM maturation site in HEK mitochondria, a semi-tryptic search was performed. For identification of unknown PTMs, the data was analyzed using the open search function of pFind 3.1.5 ^39^ with a precursor and fragment mass tolerance of 10 ppm. Consensus identification lists were generated with false discovery rates set at 1% for protein, peptide, and site identifications.

### Northern blotting

The sperm and HEK293 RNA samples were diluted in a loading buffer (95% formamide, 5 mM EDTA, 0.025% SDS, 0.025% bromophenol blue, and 0.025% xylene cyanol) and heated at 95°C for 3 minutes to further denature samples before loading. RNA (1 μg) was loaded onto a 6% PAGE containing 6M Urea in 1X Tris-Borate-EDTA (TBE) buffer. HEK293 RNA (50 ng) was loaded onto the gel, and detection of 18S rRNA was used as a size marker. The RNA species were transferred onto a Hybond-H+ membrane (GE Healthcare) in 1X TBE buffer and crosslinked by UV exposure for 2 minutes upon electrophoresis. The DNA probes (**Extended Table 1**) at 500 nM were 5’-[^32^P]-labeled using T4 Polynucleotide Kinase (NEB) and hybridized to the membrane overnight at 37°C in PerfectHyb™ Plus Hybridization Buffer (Sigma-Aldrich). The membrane was washed twice with 2X SSC buffer (Sigma-Aldrich), dried, and visualized by autoradiography using PhosphorImager (GE Healthcare).

### Western blotting

Spermatozoa (∼ 5 million cells) were lysed in 100 μl of 1x Laemmli sample buffer containing 10% 2-Mercaptoethanol by sonication. Proteins were separated using 12 %SDS-PAGE and blotted onto a polyvinylidene difluoride membrane (GE Healthcare). The membrane was blocked in 5% milk/PBS for 1 h followed by overnight incubation with mouse monoclonal anti-TFAM antibody (Abcam, #ab119684, 1:1000 dilution) at 4 °C. PolG was detected using anti-POLG antibody (Abcam, #128899, 1:1000). TEFM was detected using home-raised mouse monoclonal antibody against human TEFM (1:1000), while mtRNAP - using home-raised rabbit polyclonal antibodies against human Δ150 mtRNAP (1:1000). The membrane was washed three times with PBST (1X PBS and 0.1 % Tween) and incubated with IRDye 680RD-labeled goat anti-mouse antibody (LI-Cor, #926-68070, 1:10,000 dilution) or IRDye 800CW-labeled goat anti-Rabbit IgG Secondary Antibody (LI-Cor, #926-32211, 1:15000) for 1 h at RT. The membrane was washed 6 times with PBST and imaged using the Infrared Imaging System Odyssey FC (LI-Cor).

### Cloning

For the lentiviral expression, the DNA region encoding TFAM-mScarlet fusion was excised from the pcDNA3-TFAM-mScarlet plasmid (Addgene ref #129573) using BamHI and EcoRI endonucleases and inserted into the pWPXL plasmid (Addgene ref #12257). To obtain the TFAM construct containing the 3’mitoUTR (3’mitoUTR_TFAM-mScarlet_pWPXL), human heart cDNA (Zyagen, HD-801) was PCR-amplified using primers described in **Extended Data Table 2**, and the amplicon cloned into a pT7Blue cloning vector (Novagen). Restriction sites (EcoRI/NdeI) were added through amplification by PCR to insert the fragment into the TFAM-mScarlet/pWPXL plasmid. The 5’mitoUTR was cloned into the TFAM-mScarlet/pWPXL or the 3’mitoUTR_TFAM-mScarlet_pWPXL plasmid using megaprimers following the QuikChange II XL Site-Directed Mutagenesis Kit (Agilent) large insertion protocol provided by the manufacturer. The Megaprimers for the 5’ mitoUTR were synthesized by amplifying DNA extracted from HEK293 cells using primers (**Extended Data Table 2**). To obtain the TFAM constructs containing the 3’ and/or 5’ sperm TFAM UTRs, megaprimers were synthesized and cloned into the TFAM-mScarlet_pWPXL plasmid following the QuikChange II XL Site-Directed Mutagenesis Kit (Agilent) large insertion protocol. The Megaprimers were synthesized by amplifying DNA extracted from HEK293 (**Extended Data Table 2**). Deletion of the TFAM MTS (residues 1-42, Δ42TFAM-mScarlet) was performed using the QuikChange II XL Site-Directed Mutagenesis kit in 3’5’mitoUTR_TFAM-mScarlet_pWPXL.

TOM20 and H2B Type 1A ORFs were amplified by PCR from human heart cDNA (Zyagen). The amplicons were cloned into the pT7Blue cloning vector (Novagen) using the megaprimers (**Extended Data Table 2**). The protocol for large insertions provided by the QuikChange II XL Site-Directed Mutagenesis Kit (Agilent) was followed to insert TOM20 or H2B in place of the TFAM gene TFAM-mScarlet_pWPXL plasmid. The MTS^TFAM^mScarlet_pWPXL construct was generated using QuikChange II XL site-directed mutagenesis kit by removing the TFAM sequence (residues 43-246) from the TFAM-mScarlet_pWPXL construct described above. To substitute the TFAM MTS with the mtRNAP MTS and generate the MTS^pol^1′42TFAM-mScarlet construct, we generated megaprimers by PCR amplification of the MTS region (residues 1-45) of mtRNAP. The megaprimers were inserted into the TFAM-Scarlet_pWPXL plasmid.

Human mtRNAP (residues 1-1230) was cloned into pWPXL vector to obtain mtRNAP_pWPXL construct. Megaprimers containing the mScarlet gene were used to obtain the MTS^Pol^-mScarlet-mtRNAP-pWPXL construct. The MTS^pol^mScarlet_pWPXL construct was obtained by excising the MTS^Pol^-mScarlet region using PmeI and XcmI and ligating this fragment into the pWPXL plasmid. Substitutions of the serine residues in the TFAM MTS (S31A/S34A) were performed using QuikChange II XL Site-Directed mutagenesis kit in the TFAM-mScarlet_pWPXL construct.

### Generation of lentiviral particles

In brief, 15 μg of the vector plasmid containing the gene of interest, 10 μg of the packaging construct plasmid psPAX2 (Adgene #12260), and 5 μg of VSV-G envelope expressing plasmid pMD2.G (Addgene #12259) were co-transfected into HEK293T cells using calcium phosphate transfection. The transfection mix was removed after 16 h and replaced with the complete media. The lentiviral particles were harvested after 24 and 48 h incubation, filtered through a 0.45 mm filter and concentrated by ultracentrifugation before being stored at −80 °C. The infection efficiency of the viral particles was tested on HEK293T cells to ensure over 90% infection.

### Lentiviral transduction of human cells and spermatozoa

HeLa cells were seeded at a concentration of 415 cells/mm^2^ in a 6-well plate to reach 90% confluency over 24 h before transduction. Upon incubating the lentiviral particles with HeLa cells for 16 h, the cells were washed twice with PBS and resuspended in a fresh complete culture medium. Transduced HeLa cells were obtained after 48 h post-transduction and resuspended in a fresh culture medium to be seeded for immunofluorescence (IF) staining. Viruses generated using the pWPXL vector were used as control.

To transduce the sperm cells, lentiviral particles were added to 1 ml of spermatozoa (∼10^7^ cells) in a capacitation medium and incubated for 3-48 h. The transduced sperms were centrifuged at 600 g for 5 min, and the sperm pellet was resuspended in PBS to be further spotted on coverslips for IF staining.

### Immunofluorescence staining

Before staining, HeLa cells were seeded on poly-L-lysine (Gibco) coated glass coverslips overnight, while human spermatozoa were spotted and air-dried directly on the coverslips after isolation. The samples were fixed with 4% paraformaldehyde (Thermo Fisher Scientific) for 20 minutes at RT, washed with PBS, permeabilized in PSB solution containing 0.25% Triton X-100 (Sigma-Aldrich) for 5 minutes, and blocked with PBS solution containing 1% BSA (Sigma-Aldrich) and 0.1% Triton X-100 for 1 h at 37 °C. Followed by washing with PBS, the samples were blocked with 1% BSA-PBS solution for 1 h at 37 °C. The prepared coverslips were incubated with the following primary antibodies prepared in mouse monoclonal anti-TFAM antibody (Abcam, #ab119684, 1:500) or monoclonal anti-TOM20 antibody (Cell Signaling Technology, #42406S, 1:500) in 0.1% Triton X-100 overnight at 4 °C. The coverslips were washed three times with 0.1% Triton X-100 at 4 °C for 5 min, and the secondary antibody, donkey anti-mouse Alexa Fluor 555 (Abcam #ab150110, 1:2000) or goat anti-rabbit Oregon Green 488 (Invitrogen #O-11038, 1:2000) was added for 2 h at RT. The coverslips were then counterstained with 300 nM DAPI (Invitrogen) and mounted with Prolong Glass antifade (Invitrogen). Digital images were obtained using a Nikon A1R+ confocal microscope at 60x magnification. The images were analyzed using the Fiji processing package on ImageJ.

### Transmission Electron Microscopy

Purified human spermatozoa were collected and fixed with freshly made fixative containing 3% paraformaldehyde and 0.1% glutaraldehyde in 0.1M phosphate buffer with 4% sucrose (pH 7.2). After washing with 0.1M phosphate buffer with 4% sucrose, 0.1M glycine was added to quench the unbonded aldehyde group. The cells were dehydrated in graded series of ethanol on ice and embedded in LR White (Electron Microscopy Sciences, Hatfield, PA). The samples were polymerized under UV light (360nm) at −10°C for 48 h, followed by 12 h at RT. The 90 nm-thin sections were cut and mounted on Formvar-Carbon coated 200 mesh nickel grids. The grid was incubated with an anti-TFAM antibody (1:1000, Abcam, #ab119684) in PBS with 1% BSA, and 0.05% Tween 20, for 2 h at RT, then overnight at 4°C. Upon washing, the anti-mouse secondary antibody conjugated with 18 nm gold (1:15, Colloidal Gold AffiniPure Goat Anti-Mouse IgG (H+L, EM Grade, Jackson ImmunoReasearch Laboratories, Inc.,) were added in PBS with 1% BSA, and 0.05% Tween 20, and incubated for 1h. After washing, the grids were stained with uranyl acetate and lead citrate by standard methods, imaged with Talos120C transmission electron microscope (Thermo Fisher Scientific), and recorded using Gatan (4k x 4k) OneView Camera with software Digital Micrograph (Gatan Inc).

## Contributions

Tissue collection: NMG, LL

Cloning: GB, WL, AZO

Droplet digital PCR: PP, MP, AC, RT

Protein purification: AZO

Cell culture, transduction, IH, CISH: WL

LC MS/MS analysis: HYT, WL

Regulatory Oversight: AK

Experimental design: WL, AZO, GB, RT, SM, DT,

Writing – original draft: DT

Writing – review & editing: DT, WL, AZO, RT, MA, SM

## Acknowledgments

We thank NYULH DART Microscopy Laboratory, Alice Liang, Chris Petzold, and Kristen Dancel-Manning for consultation and assistance with TEM work; this core is partially funded by NYU Cancer Center Support Grant NIH/NCI P30CA016087. The authors are indebted to Crystal Van Dyken, David Battaglia, and the staff of the Oregon National Primate Research Center, OHSU Reproductive Endocrinology, and IVF clinic for their expertise and services in obtaining monkey and human gametes for this study. We thank Thomas Jefferson University BioImaging facility and Dr. Maria Covarrubias for help with confocal microscopy experiments. Dr Michael Anikin and William T. McAllister are acknowledged for the critical reading of the manuscript and fruitful discussion.

## Funding

National Institutes of Health grant R35 GM131832 (DT)

PID2020-115091RB-I00, MCIN/AEI/10.13039/501100011033 Spain (RT)

PI2020/09-4, CIBERNED, Instituto de Salud Carlos III (ISCIII) Spain (RT)

## Extended Data for

**Extended Data Fig. 1.**
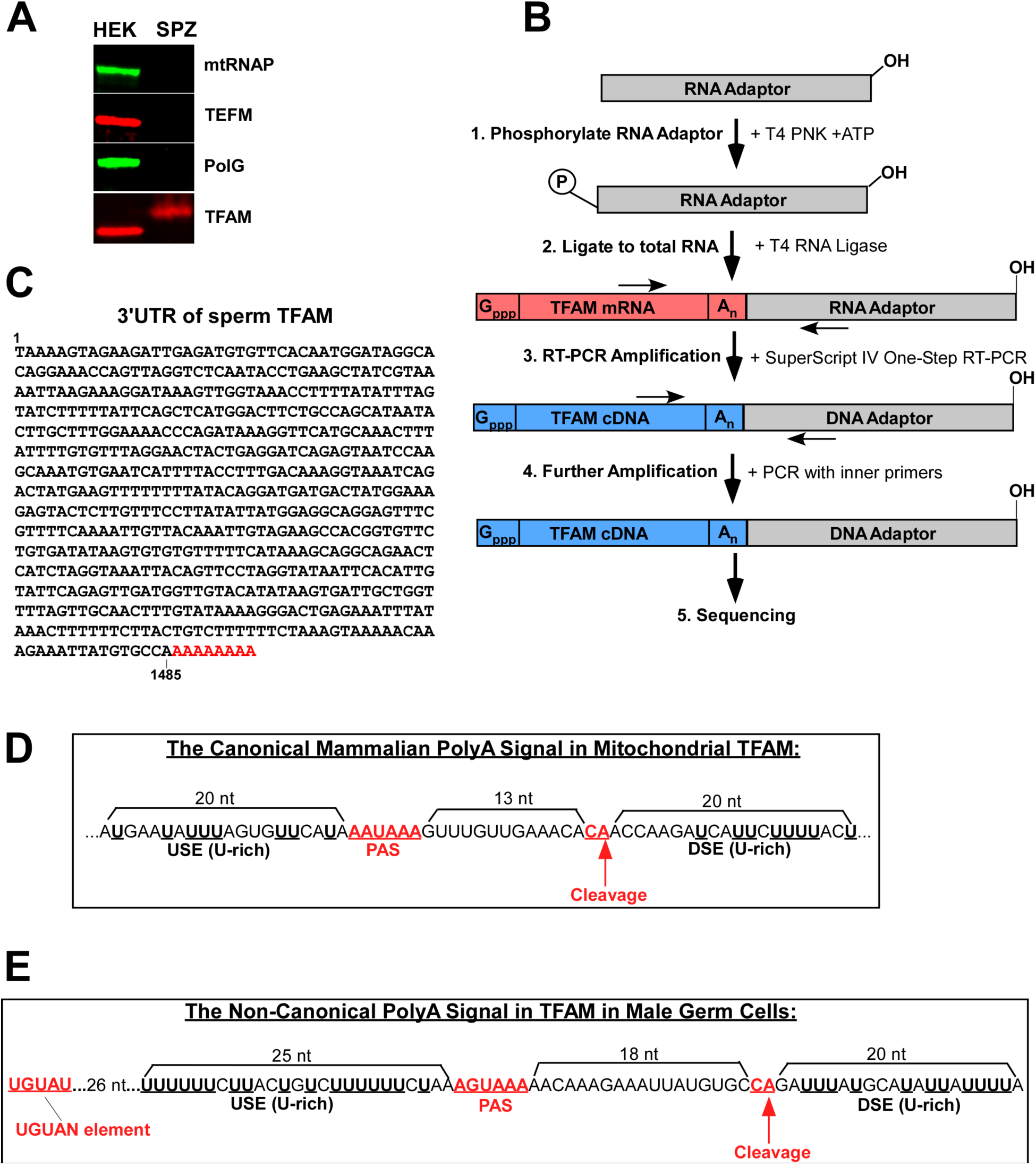
Mapping of the UTRs of the sperm TFAM isoform. **A.** Western blot analysis of the proteins indicated in mitochondria of HEK cells and spermatozoa (SPZ). **B.** Schematics of the RACE experiment. **C.** The sequence of the 3’ UTR of the sperm TFAM cDNA **D.** Schematic illustration of polyadenylation of the TFAM mRNA in somatic cells. PAS – polyadenylation signal. **E.** Schematic illustration of polyadenylation of the TFAM mRNA in sperm cells. Putative polyadenylation signal (PAS) and UGUAN elements are indicated.

**Extended Data Fig. 2.**
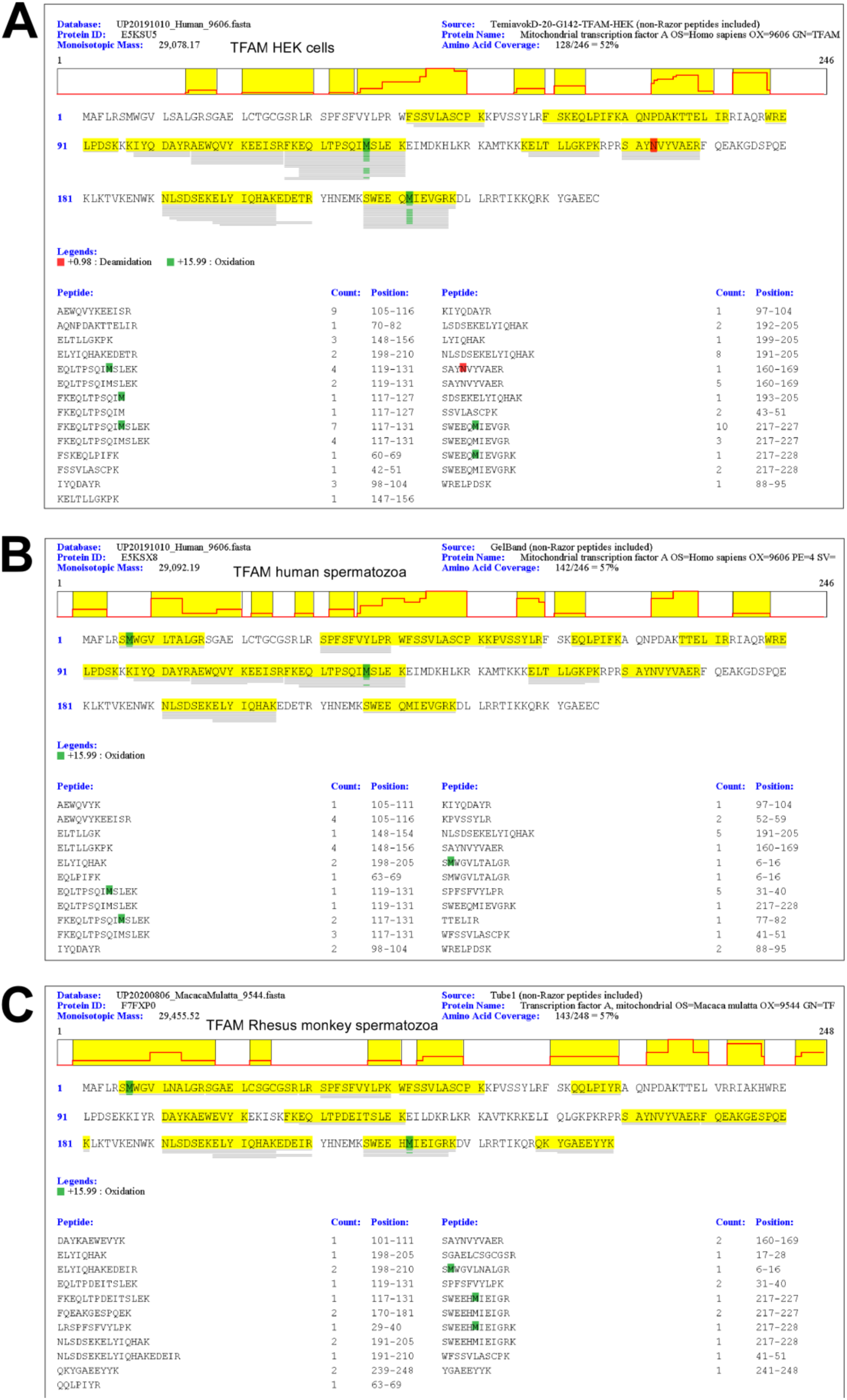
Identification of TFAM peptides by LC-MS/MS analysis in HEK cells (A), human (B) and Rhesusmonkey (C) spermatozoa. Peptide search was done using MaxQuant. Note that the SGAELCSGCGSR peptide was identified in the human sperm TFAM sample using pFind and therefore not shown in panel B.

**Extended Data Fig. 3.**
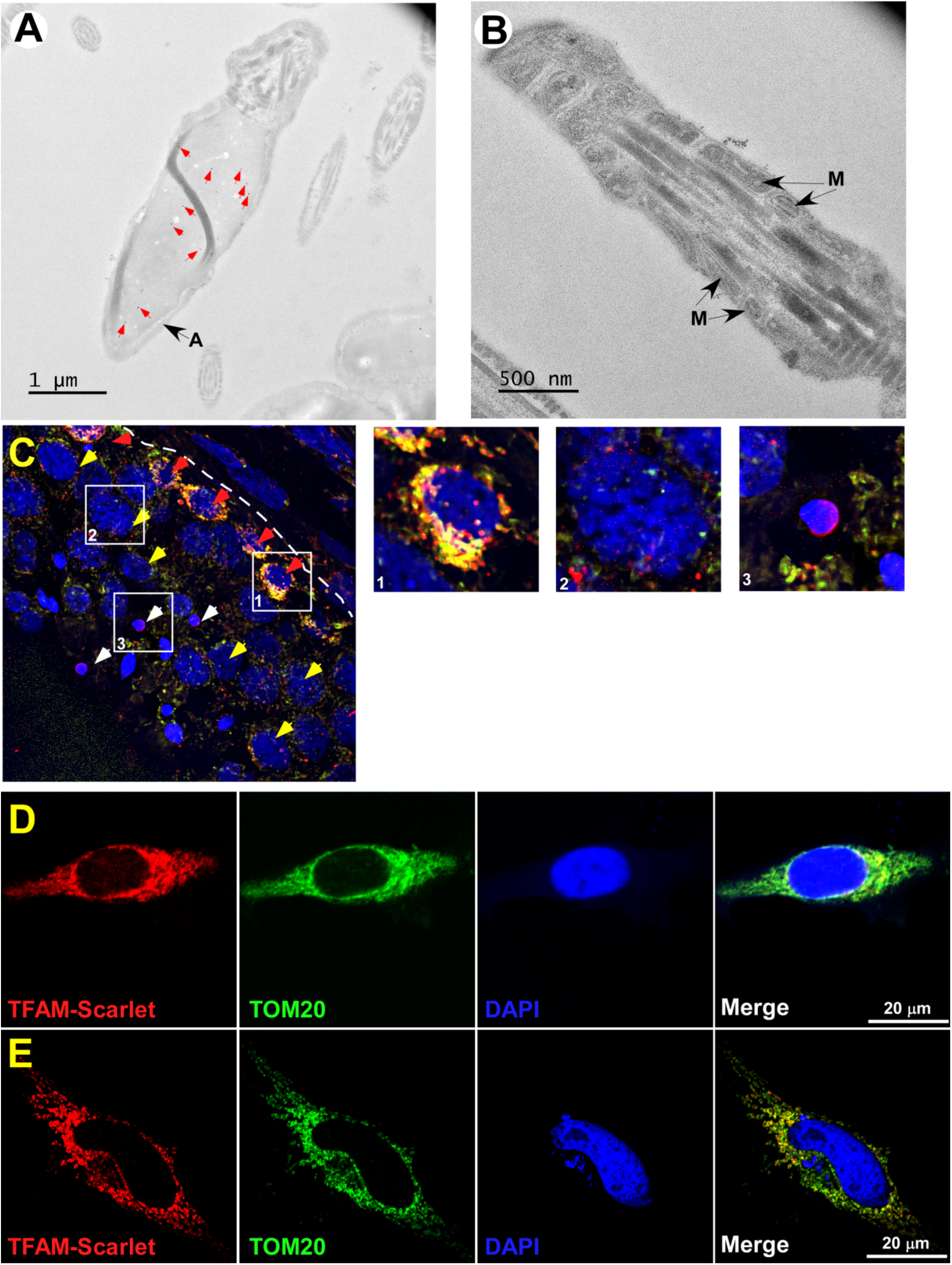
TFAM is localization in spermatozoa and somatic cells. **A.** Cryo immunogold electron microscopy of spermatozoa head. Red arrows indicate gold particles. A -acrosome. **B.** Cryo immunogold electron microscopy of spermatozoon midpiece. M- mitochondria. **C.** Staining of human testicular tissue. Merge image. Red- staining with anti-TFAM antibody, blue- DAPI, green - staining with anti-TOM20 antibody. Red arrows - spermatogonia, yellow – spermatocytes, white – spermatids. The basement membrane is indicated by a dashed line, L- seminiferous tubule lumen. Close-up images of spermatogonia (1), spermatocytes (2) and spermatids (3) correspond to the white squares indicated. **D.** Over-expressed TFAM having the somatic 5’ and 3’ UTRs shows mitochondrial localization in HeLa cells. **E.** Overexpressed TFAM lacking UTRs shows mitochondrial localization in HeLa cells

**Extended Data Fig. 4.**
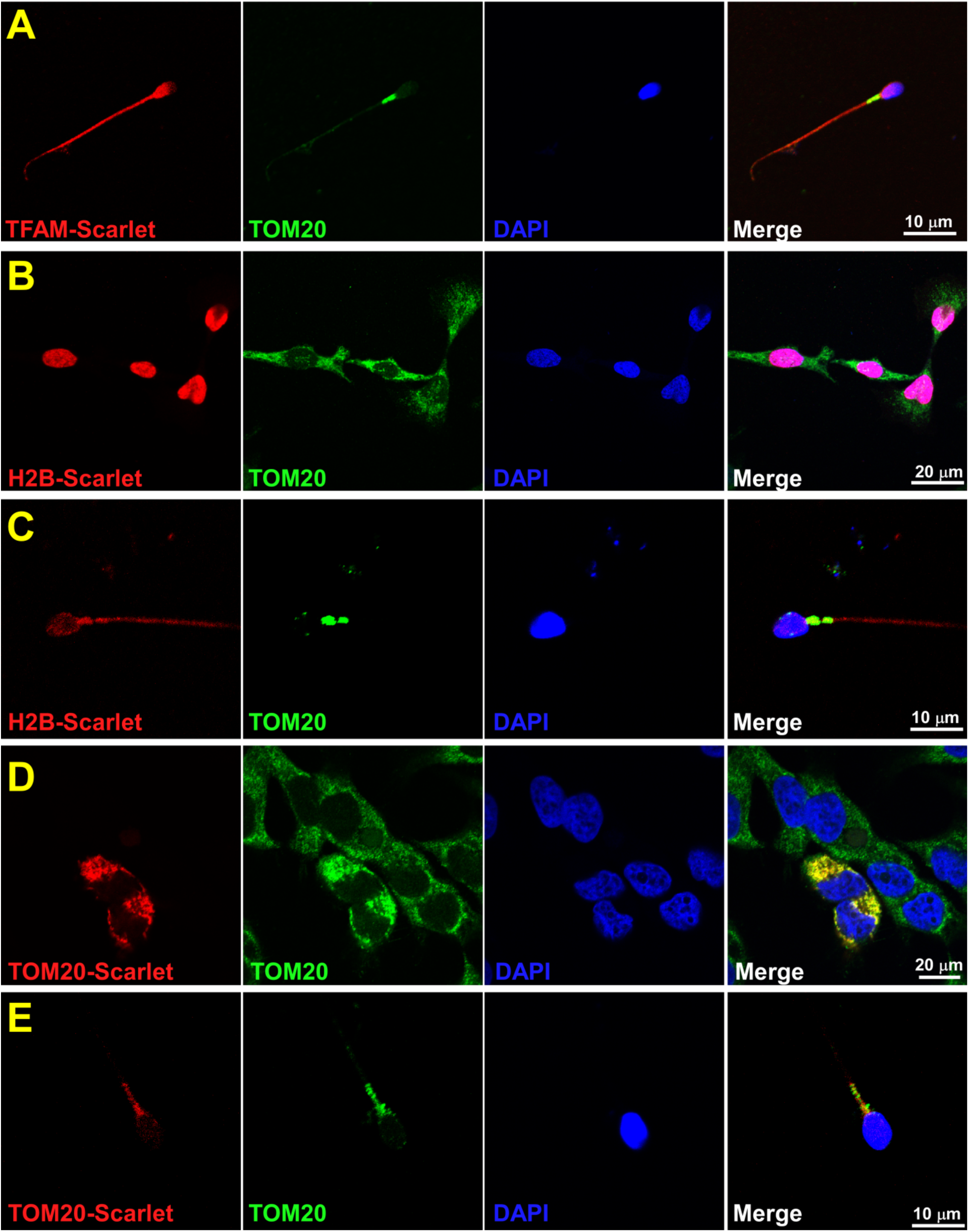
Expression of TFAM, H2B, and TOM20 in somatic cells and mature spermatozoa. **A.** Over-expression of TFAM mRNA having sperm 3’ and 5’ UTR regions result in cytoplasmic localization of this protein in spermatozoa. **B.** Over-expression of H2B in HeLa cells shows nuclear localization. **C.** Over-expression of H2B in sperm cells results in cytoplasmic localization. **D.** Over-expression of TOM20 shows mitochondrial localization in HeLa cells. **E.** Over-expression of TOM20 results in mitochondrial localization in sperm cells

**Extended Data Fig. 5.**
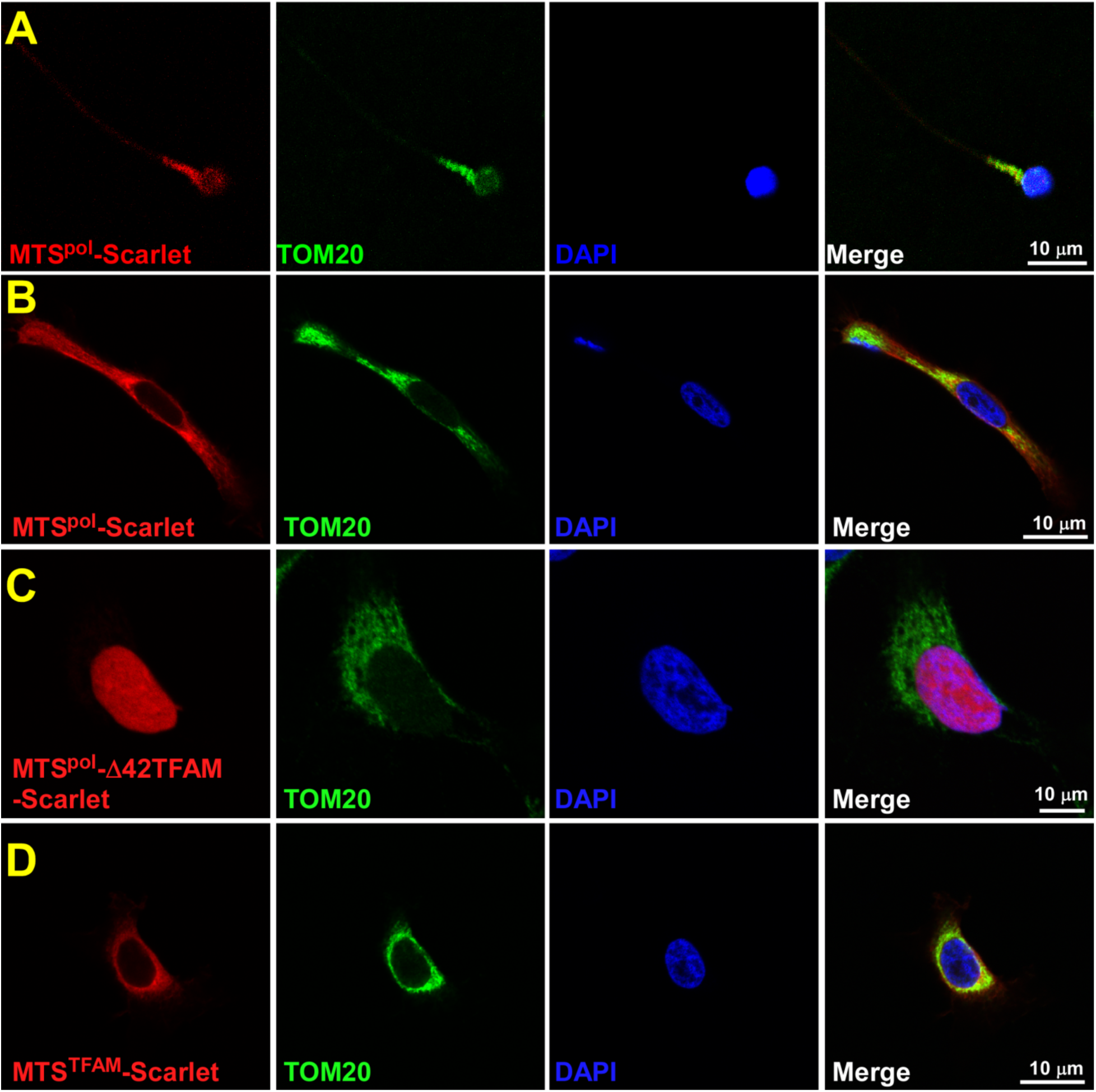
Trafficking of the TFAM variants in HeLa cells. **A.** MTS_pol_-mScarlet protein is localized to the sperm mitochondria. **B.** MTS_pol_-mScarlet is localized to mitochondria in HeLa cells. **C.** MTS_Pol_-D42TFAM is localized to the nucleus of HeLa cells. **D.** MTS_TFAM_-Scarlet is localized to mitochondria in HeLa cells

**Extended Data Table 1.**
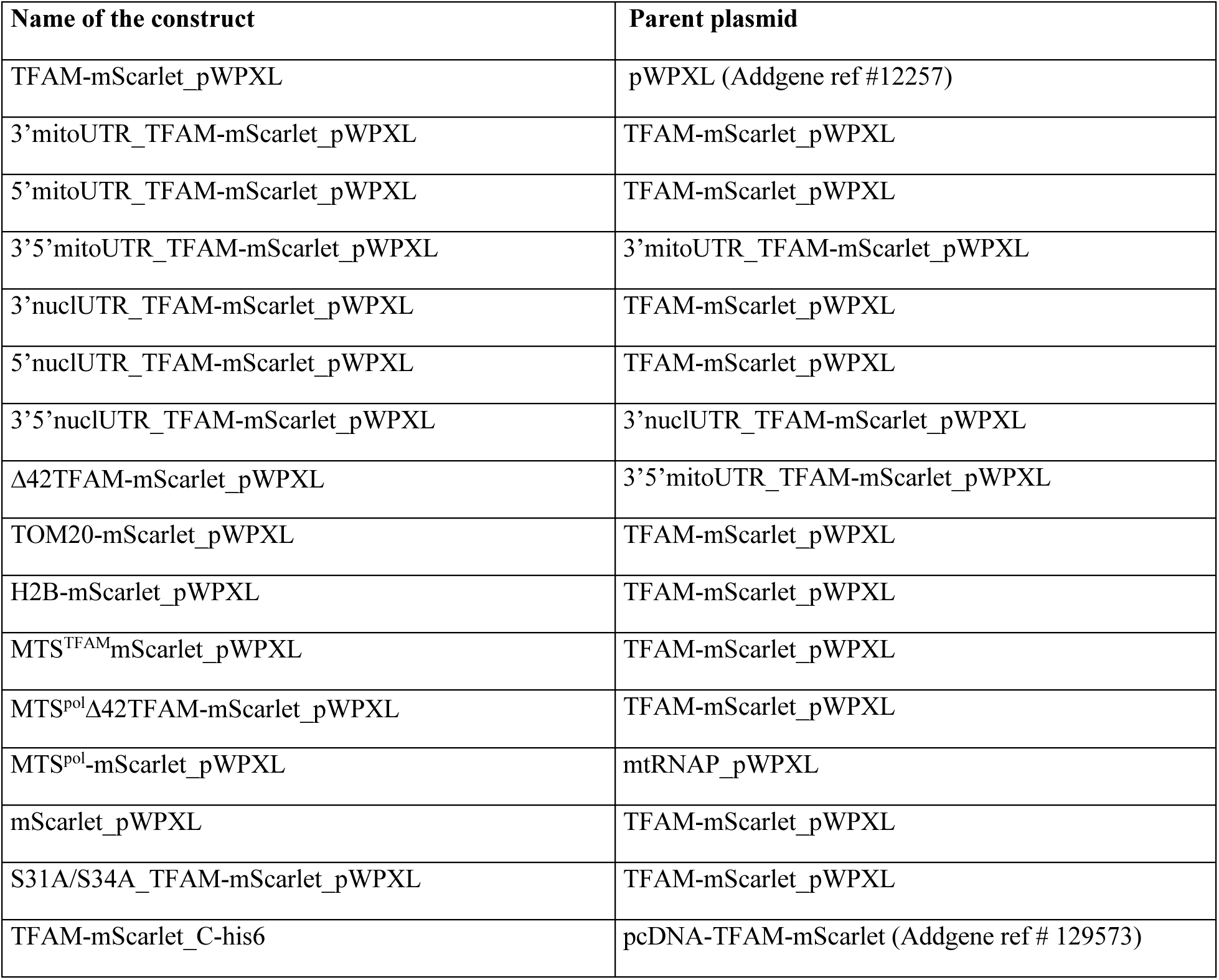
Plasmids generated during the study.

**Extended Data Table 2.**
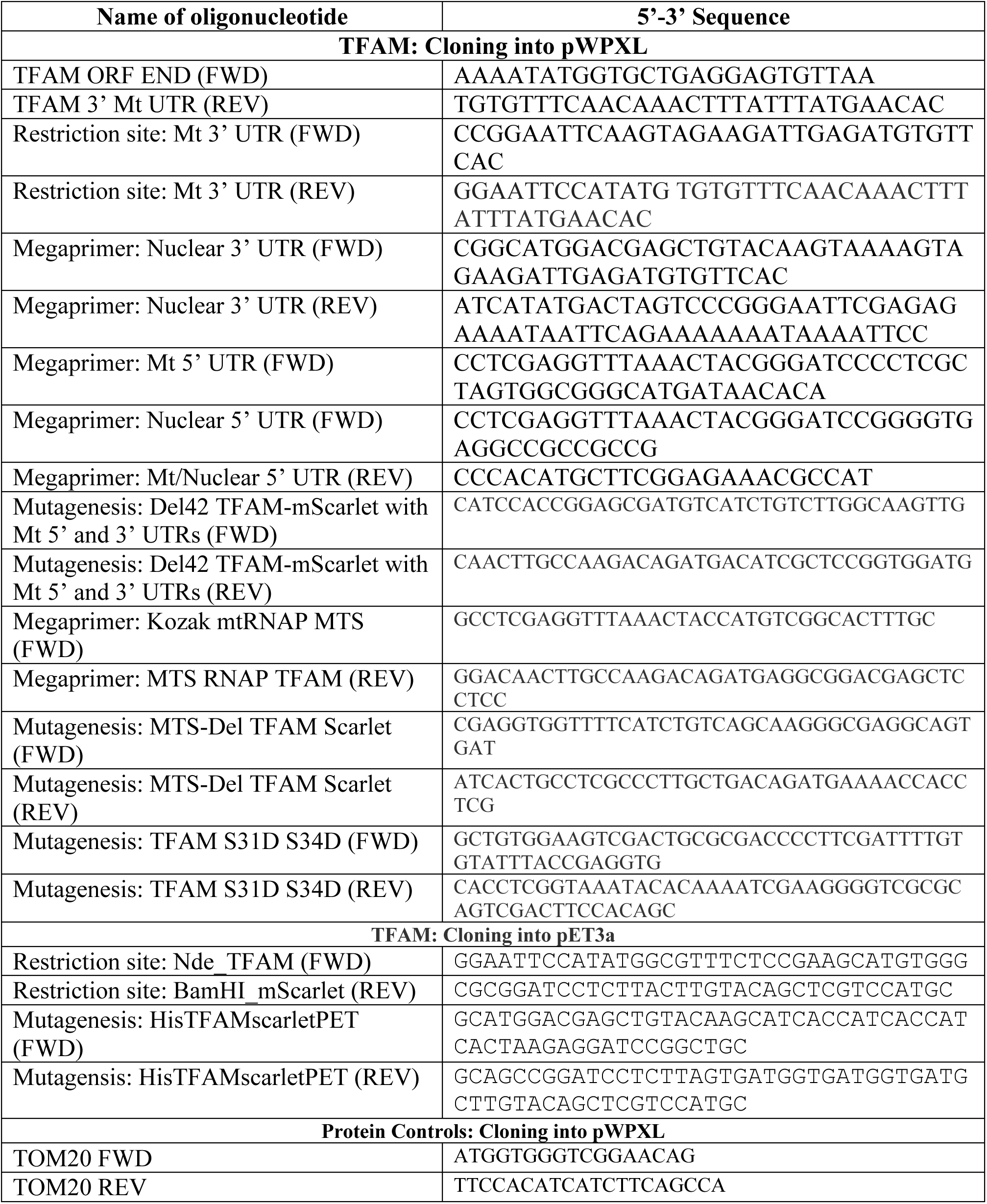
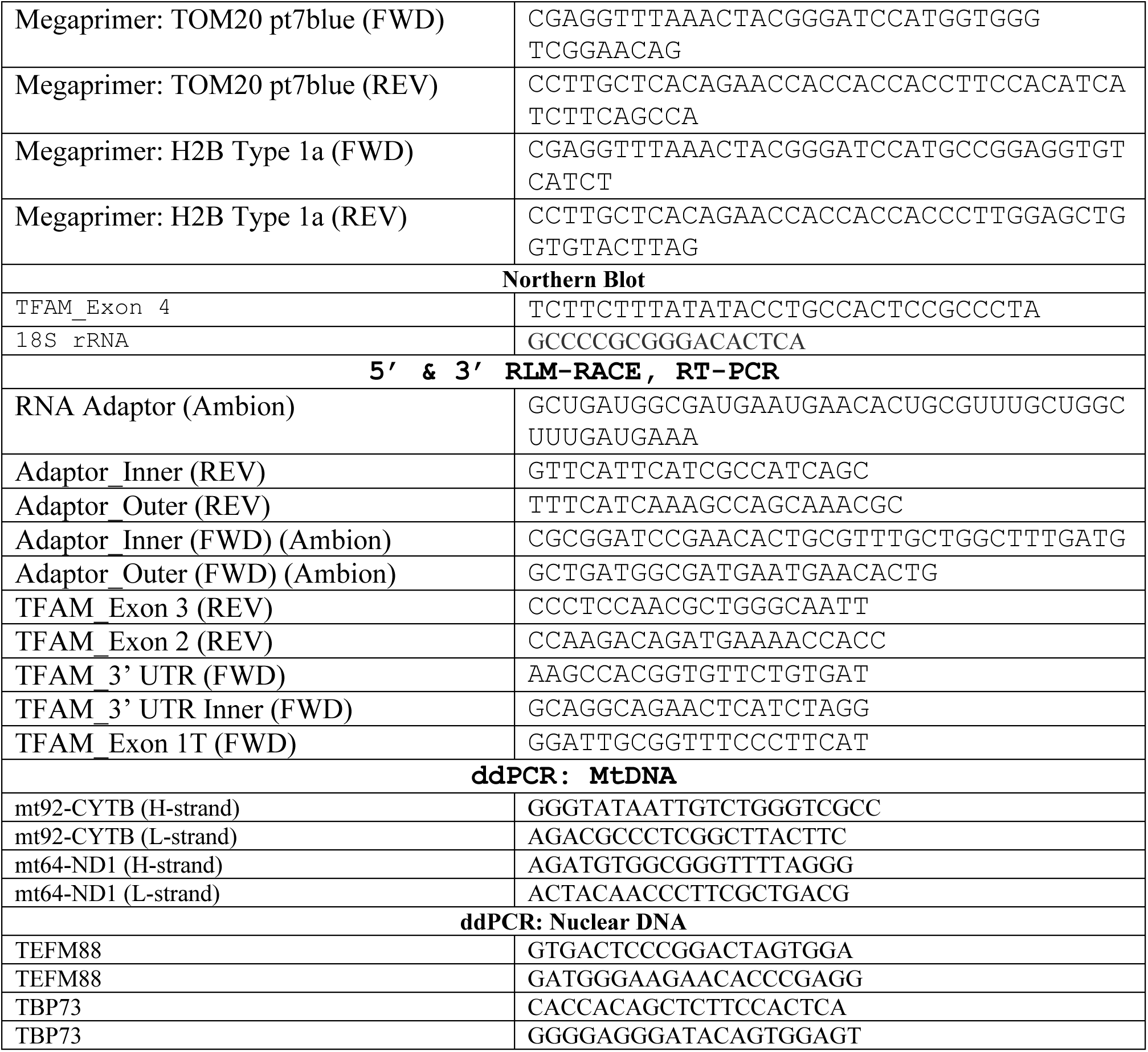
Oligonucleotides used in the study.

