## Supplemental Figures for "Molecular Basis for Maternal Inheritance of Human Mitochondrial DNA"

[1] Department of Biochemistry, Thomas Jefferson University; 1020 Locust St, Philadelphia 19107, USA, [2]Neurobiology Unit, Institut d'Investigacions Biomèdiques de Barcelona (IIBB-CSIC-IDIBAPS) and Centro de Investigación Biomédica en Red de Enfermedades Neurodegenerativas (CIBERNED); Barcelona, 08036, Spain, [3]Center for Embryonic Cell and Gene Therapy, Oregon Health & Science University; 3303 W Bond Avenue, Portland, Oregon 97239, USA, [4]The Wistar Institute, Philadelphia, 3601 Spruce Street, Philadelphia, PA 19104, USA, [5]Department of Pathology, Thomas Jefferson University; 1020 Locust St, Philadelphia 19107, USA,

**The file includes Extended data Figures 1-5 and Extended data Tables 1, 2**

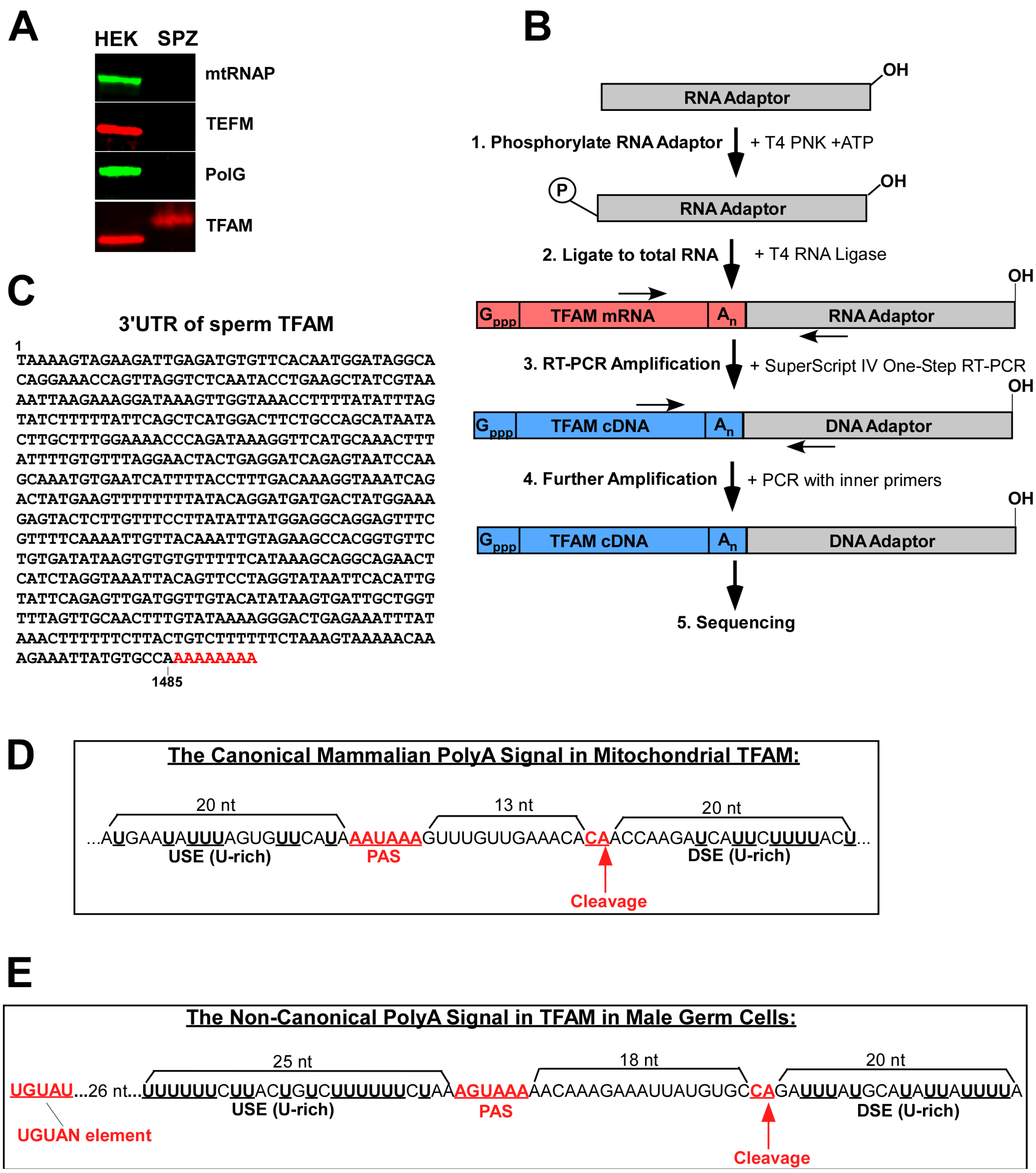


**Extended Data Fig. 1. Mapping of the UTRs of the sperm TFAM isoform.**
**A.** Western blot analysis of the proteins indicated in mitochondria of HEK cells and spermatozoa (SPZ).
**B.** Schematics of the RACE experiment.
**C.** The sequence of the 3’ UTR of the sperm TFAM cDNA
**D.** Schematic illustration of polyadenylation of the TFAM mRNA in somatic cells. PAS – polyadenylation signal.
**E.** Schematic illustration of polyadenylation of the TFAM mRNA in sperm cells. Putative polyadenylation signal (PAS) and UGUAN elements are indicated.

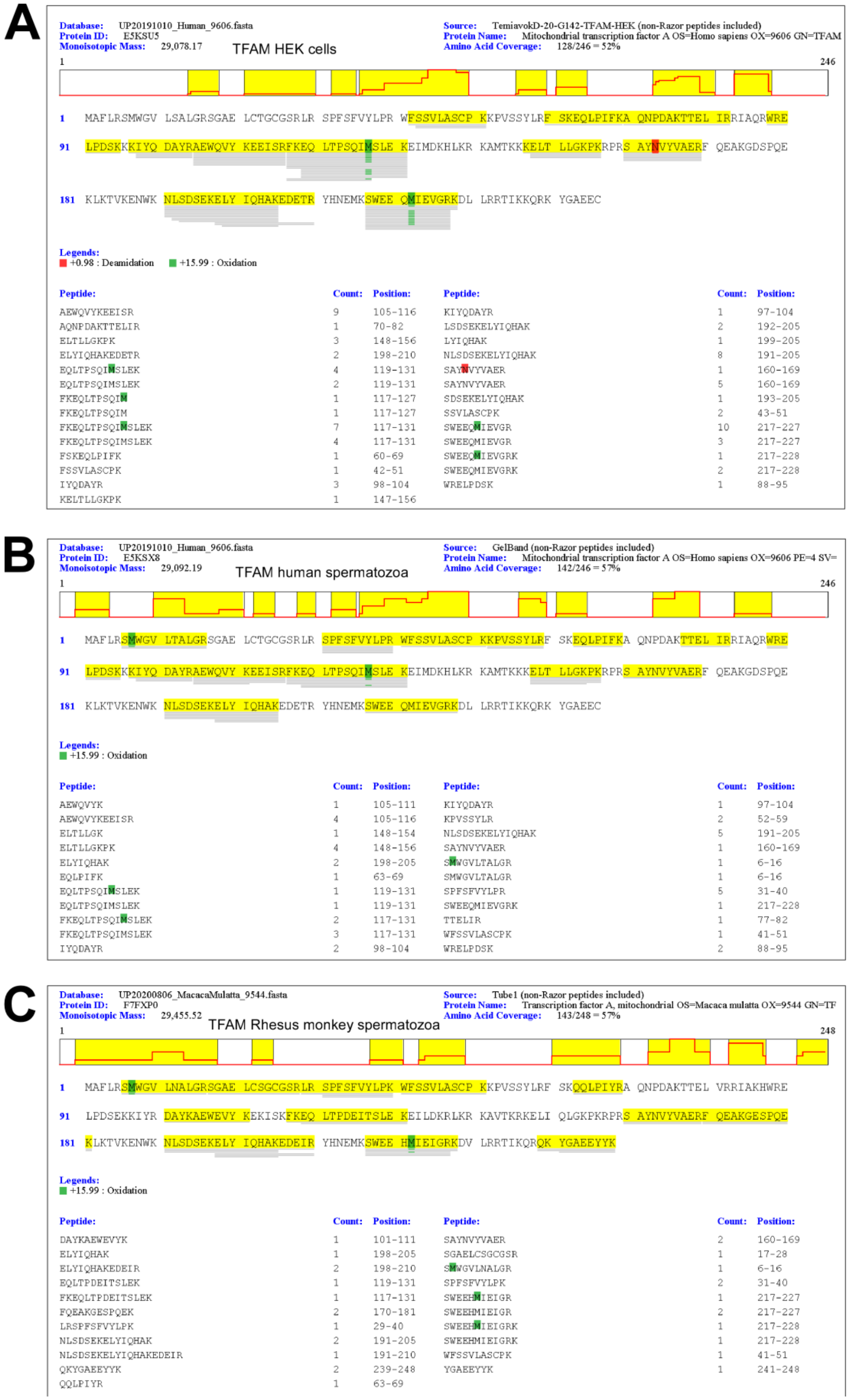


**Extended Data Fig.2**. **Identification of TFAM peptides by LC-MS/MS analysis in HEK cells (A), human (B) and Rhesusmonkey (C) spermatozoa**. Peptide search was done using MaxQuant. Note that the SGAELCSGCGSR peptide was identified in the human sperm TFAM sample using pFind and therefore not shown in panel B.

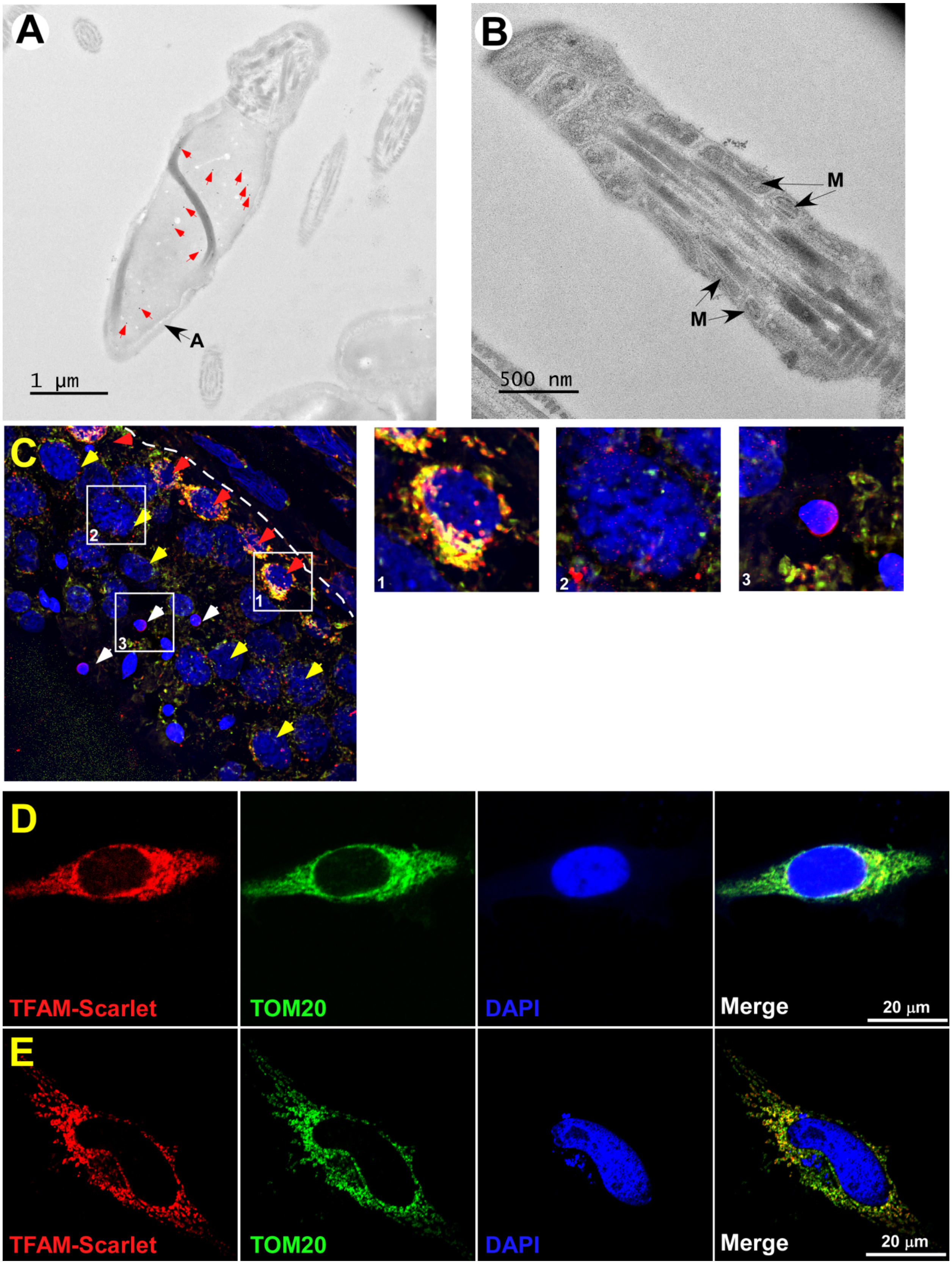


**Extended Data Fig. 3. TFAM is localization in spermatozoa and somatic cells.**

**A.** Cryo immunogold electron microscopy of spermatozoa head. Red arrows indicate gold particles. A -acrosome.
**B**. Cryo immunogold electron microscopy of spermatozoon midpiece. M- mitochondria.
**C.** Staining of human testicular tissue. Merge image. Red- staining with anti-TFAM antibody, blue- DAPI, green - staining with anti-TOM20 antibody. Red arrows - spermatogonia, yellow – spermatocytes, white – spermatids. The basement membrane is indicated by a dashed line, L- seminiferous tubule lumen. Close-up images of spermatogonia (1), spermatocytes (2) and spermatids (3) correspond to the white squares indicated.
**D.** Over-expressed TFAM having the somatic 5' and 3' UTRs shows mitochondrial localization in HeLa cells.
**E.** Overexpressed TFAM lacking UTRs shows mitochondrial localization in HeLa cells

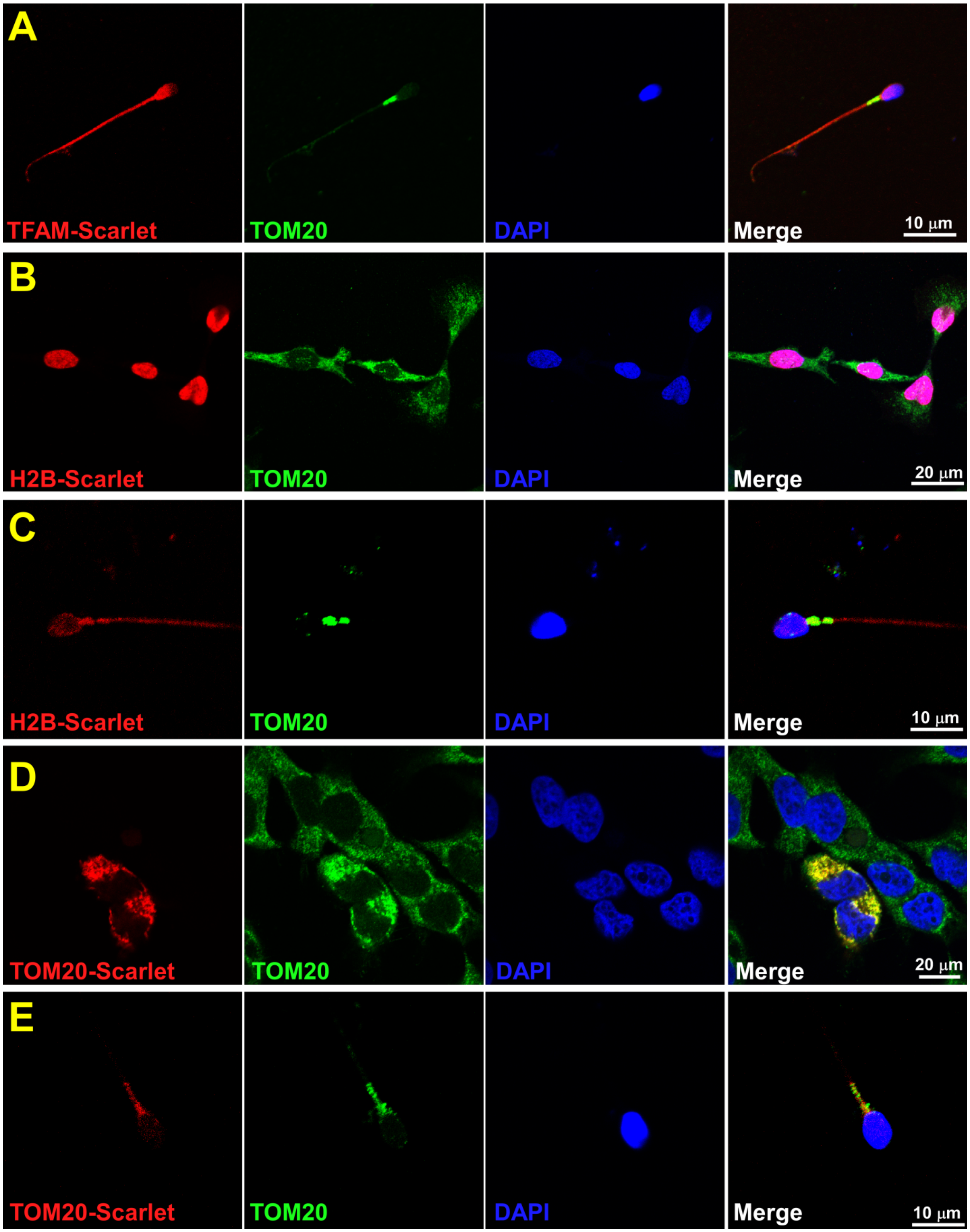


**Extended Data Fig.4. Expression of TFAM, H2B, and TOM20 in somatic cells and mature spermatozoa.**

**A.** Over-expression of TFAM mRNA having sperm 3’ and 5’ UTR regions result in cytoplasmic localization of this protein in spermatozoa.
**B.** Over-expression of H2B in HeLa cells shows nuclear localization.
**C.** Over-expression of H2B in sperm cells results in cytoplasmic localization.
**D.** Over-expression of TOM20 shows mitochondrial localization in HeLa cells.
**E.** Over-expression of TOM20 results in mitochondrial localization in sperm cells

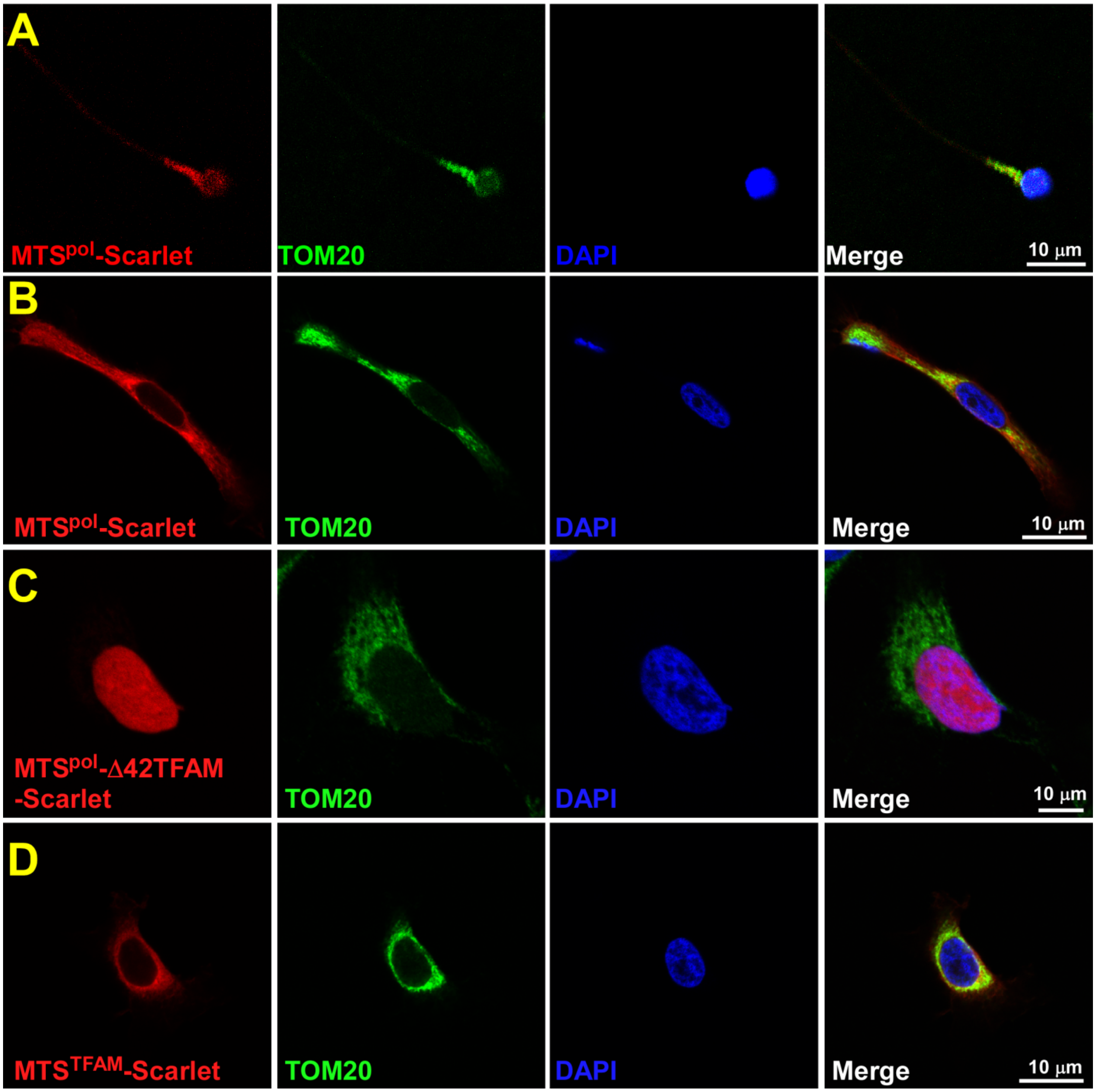


**Extended Data Fig. 5. Trafficking of the TFAM variants in HeLa cells.**
**A.** MTS$_{pol}$-mScarlet protein is localized to the sperm mitochondria.
**B.** MTS$_{pol}$-mScarlet is localized to mitochondria in HeLa cells.
**C.** MTS$_{Pol}$-D42TFAM is localized to the nucleus of HeLa cells.
**D.** MTS$_{TFAM}$-Scarlet is localized to mitochondria in HeLa cells

**Extended Data Table 1. Plasmids generated during the study**

| **Name of the construct** | **Parent plasmid** |
|---|---|
| TFAM-mScarlet_pWPXL | pWPXL (Addgene ref #12257) |
| 3’mitoUTR_TFAM-mScarlet_pWPXL | TFAM-mScarlet_pWPXL |
| 5’mitoUTR_TFAM-mScarlet_pWPXL | TFAM-mScarlet_pWPXL |
| 3’5’mitoUTR_TFAM-mScarlet_pWPXL | 3’mitoUTR_TFAM-mScarlet_pWPXL |
| 3’nuclUTR_TFAM-mScarlet_pWPXL | TFAM-mScarlet_pWPXL |
| 5’nuclUTR_TFAM-mScarlet_pWPXL | TFAM-mScarlet_pWPXL |
| 3’5’nuclUTR_TFAM-mScarlet_pWPXL | 3’nuclUTR_TFAM-mScarlet_pWPXL |
| Δ42TFAM-mScarlet_pWPXL | 3’5’mitoUTR_TFAM-mScarlet_pWPXL |
| TOM20-mScarlet_pWPXL | TFAM-mScarlet_pWPXL |
| H2B-mScarlet_pWPXL | TFAM-mScarlet_pWPXL |
| $MTS^{TFAM}$mScarlet_pWPXL | TFAM-mScarlet_pWPXL |
| $MTS^{pol}$Δ42TFAM-mScarlet_pWPXL | TFAM-mScarlet_pWPXL |
| $MTS^{pol}$-mScarlet_pWPXL | mtRNAP_pWPXL |
| mScarlet_pWPXL | TFAM-mScarlet_pWPXL |
| S31A/S34A_TFAM-mScarlet_pWPXL | TFAM-mScarlet_pWPXL |
| TFAM-mScarlet_C-his6 | pcDNA-TFAM-mScarlet (Addgene ref # 129573) |

**Extended Data Table 2. Oligonucleotides used in the study**

| Name of oligonucleotide | 5'-3' Sequence |
|---|---|
| **TFAM: Cloning into pWPXL** | |
| TFAM ORF END (FWD) | AAAATATGGTGCTGAGGAGTGTTAA |
| TFAM 3' Mt UTR (REV) | TGTGTTTCAACAAACTTTATTTATGAACAC |
| Restriction site: Mt 3' UTR (FWD) | CCGGAATTCAAGTAGAAGATTGAGATGTGTTCAC |
| Restriction site: Mt 3' UTR (REV) | GGAATTCCATATG TGTGTTTCAACAAACTTTATTTATGAACAC |
| Megaprimer: Nuclear 3' UTR (FWD) | CGGCATGGACGAGCTGTACAAGTAAAAGTAGAAGATTGAGATGTGTTCAC |
| Megaprimer: Nuclear 3' UTR (REV) | ATCATATGACTAGTCCCGGGAATTCGAGAGAAAATAATTCAGAAAAAAATAAAATTCC |
| Megaprimer: Mt 5' UTR (FWD) | CCTCGAGGTTTAAACTACGGGATCCCCTCGCTAGTGGCGGGCATGATAACACA |
| Megaprimer: Nuclear 5' UTR (FWD) | CCTCGAGGTTTAAACTACGGGATCCGGGGTGAGGCCGCCGCCG |
| Megaprimer: Mt/Nuclear 5' UTR (REV) | CCCACATGCTTCGGAGAAACGCCAT |
| Mutagenesis: Del42 TFAM-mScarlet with Mt 5' and 3' UTRs (FWD) | CATCCACCGGAGCGATGTCATCTGTCTTGGCAAGTTG |
| Mutagenesis: Del42 TFAM-mScarlet with Mt 5' and 3' UTRs (REV) | CAACTTGCCAAGACAGATGACATCGCTCCGGTGGATG |
| Megaprimer: Kozak mtRNAP MTS (FWD) | GCCTCGAGGTTTAAACTACCATGTCGGCACTTTGC |
| Megaprimer: MTS RNAP TFAM (REV) | GGACAACTTGCCAAGACAGATGAGGCGGACGAGCTCCTCC |
| Mutagenesis: MTS-Del TFAM Scarlet (FWD) | CGAGGTGGTTTTCATCTGTCAGCAAGGGCGAGGCAGTGAT |
| Mutagenesis: MTS-Del TFAM Scarlet (REV) | ATCACTGCCTCGCCCTTGCTGACAGATGAAAACCACCTCG |
| Mutagenesis: TFAM S31D S34D (FWD) | GCTGTGGAAGTCGACTGCGCGACCCCTTCGATTTTGTGTATTTACCGAGGTG |
| Mutagenesis: TFAM S31D S34D (REV) | CACCTCGGTAAATACACAAAATCGAAGGGGTCGCGCAGTCGACTTCCACAGC |
| **TFAM: Cloning into pET3a** | |
| Restriction site: Nde_TFAM (FWD) | GGAATTCCATATGGCGTTTCTCCGAAGCATGTGGG |
| Restriction site: BamHI_mScarlet (REV) | CGCGGATCCTCTTACTTGTACAGCTCGTCCATGC |
| Mutagenesis: HisTFAMscarletPET (FWD) | GCATGGACGAGCTGTACAAGCATCACCATCACCATCACTAAGAGGATCCGGCTGC |
| Mutagensis: HisTFAMscarletPET (REV) | GCAGCCGGATCCTCTTAGTGATGGTGATGGTGATGCTTGTACAGCTCGTCCATGC |
| **Protein Controls: Cloning into pWPXL** | |
| TOM20 FWD | ATGGTGGGTCGGAACAG |
| TOM20 REV | TTCCACATCATCTTCAGCCA |

| | |
|---|---|
| Megaprimer: TOM20 pt7blue (FWD) | CGAGGTTTAAACTACGGGATCCATGGTGGG TCGGAACAG |
| Megaprimer: TOM20 pt7blue (REV) | CCTTGCTCACAGAACCACCACCACCTTCCACATCA TCTTCAGCCA |
| Megaprimer: H2B Type 1a (FWD) | CGAGGTTTAAACTACGGGATCCATGCCGGAGGTGT CATCT |
| Megaprimer: H2B Type 1a (REV) | CCTTGCTCACAGAACCACCACCACCCTTGGAGCTG GTGTACTTAG |
| **Northern Blot** | |
| TFAM_Exon 4 | TCTTCTTTATATACCTGCCACTCCGCCCTA |
| 18S rRNA | GCCCCGCGGGACACTCA |
| **5′ & 3′ RLM-RACE, RT-PCR** | |
| RNA Adaptor (Ambion) | GCUGAUGGCGAUGAAUGAACACUGCGUUUGCUGGC UUUGAUGAAA |
| Adaptor_Inner (REV) | GTTCATTCATCGCCATCAGC |
| Adaptor_Outer (REV) | TTTCATCAAAGCCAGCAAACGC |
| Adaptor_Inner (FWD) (Ambion) | CGCGGATCCGAACACTGCGTTTGCTGGCTTTGATG |
| Adaptor_Outer (FWD) (Ambion) | GCTGATGGCGATGAATGAACACTG |
| TFAM_Exon 3 (REV) | CCCTCCAACGCTGGGCAATT |
| TFAM_Exon 2 (REV) | CCAAGACAGATGAAAACCACC |
| TFAM_3' UTR (FWD) | AAGCCACGGTGTTCTGTGAT |
| TFAM_3' UTR Inner (FWD) | GCAGGCAGAACTCATCTAGG |
| TFAM_Exon 1T (FWD) | GGATTGCGGTTTCCCTTCAT |
| **ddPCR: MtDNA** | |
| mt92-CYTB (H-strand) | GGGTATAATTGTCTGGGTCGCC |
| mt92-CYTB (L-strand) | AGACGCCCTCGGCTTACTTC |
| mt64-ND1 (H-strand) | AGATGTGGCGGGTTTTAGGG |
| mt64-ND1 (L-strand) | ACTACAACCCTTCGCTGACG |
| **ddPCR: Nuclear DNA** | |
| TEFM88 | GTGACTCCCGGACTAGTGGA |
| TEFM88 | GATGGGAAGAACACCCGAGG |
| TBP73 | CACCACAGCTCTTCCACTCA |
| TBP73 | GGGGAGGGATACAGTGGAGT |
